# Remodeling of self-assembled microvascular networks under long term flow

**DOI:** 10.1101/2025.03.17.643791

**Authors:** Marie Floryan, Elena Cambria, Adriana Blazeski, Mark F. Coughlin, Zhengpeng Wan, Giovanni Offeddu, Vinayak Vinayak, Aayush Kant, Vivek Shenoy, Roger D. Kamm

## Abstract

The incorporation of a functional perfusable microvascular network (MVN) is a common requirement for most organ on-chip-models. Long-term perfusion of MVNs is often required for the maturation of organ phenotypes and disease pathologies and to model the transport of cells and drugs entering organs. In our microphysiological system, we observe that flow can recover perfusion in regressed MVNs and maintain perfusable MVNs for at least 51 days. Throughout the 51 days, however, the MVNs are continuously remodeling to align with the direction of bulk flow and only appear to attain morphological homeostasis with the use of maintenance medium without growth factors. We observed that the flow resistance of the MVNs decreases over time, and using a computational model, we show that stable vessels have higher flow rates and velocities compared to regressing vessels. Cytokine analysis suggests that static conditions generate an inflammatory state, and that continuous flow reduces inflammation over an extended period. Finally, through bulk RNA sequencing we identify that both the endothelial and fibroblast cells are actively engaged in vascular and matrix remodeling due to flow and that these effects persist for at least 2 weeks. This MPS can be applied to study hemodynamically driven processes, such as metastatic dissemination or drug distribution, or to model long-term diseases previously not captured by MPS, such as chronic inflammation or aging-associated diseases.

## Introduction

Incorporating continuously perfused 3D microvascular networks (MVNs) within microphysiological systems (MPS) is critical to developing novel models of disease. While the capabilities of MPS in modeling diseases have significantly expanded in recent years, there is a need to develop new vascularized MPS that allow for continuous monitoring of hemodynamic processes such as drug transport or cancer metastatic dissemination, and that capture long-term disease dynamics central to age-related cellular changes, chronic diseases, or inflammatory processes. In each case, the value of the MPS is enhanced by closely mimicking physiological flow conditions and the morphology of the organ’s specific microvascular bed. Local changes in blood flow dynamics, dictated by changing vascular morphology, are often associated with disease states, such as inflammation^1^. In the case of cancer, the inflammatory state of the endothelium is crucial as it influences immune or tumor cell adhesion, their retention at the site of adhesion, and their transendothelial migration. Furthermore, inflammation may also affect MVNs. Inflammation promotes angiogenesis, EC proliferation and migration, increases vascular permeability, and facilitates extracellular matrix (ECM) remodeling^2,3^. Capturing the inflammatory state of the endothelium is therefore critical for modeling the immune response. Continuous perfusion of MVNs is a promising approach to meet the requirements of capturing hemodynamic processes and long-term disease dynamics.

MPS that incorporate self-assembled MVNs are an attractive approach to modeling such dynamics because their morphology closely mimics that of the *in vivo* microcirculation. Such MVNs typically form over 4-7 days and are grown statically for up to 14 days total before regression sets in^4,5^. This culture period has allowed for numerous important studies: circulating immune cells have been introduced to investigate the role they play in various diseases, notably metastatic cancer^6–8^, and the subcutaneous absorption of drugs has been characterized^7^. These studies, however, lack the incorporation of hemodynamics.

Most MPS studies incorporating continuous perfusion of MVNs have limited their perfusion periods to 24-hours to 7 days (up to 14 days total) ^9–11^. Luminal flow conditioning over two days was shown to have anti-inflammatory effects^10^, and in a similar microfluidic device, luminal flow was shown to promote the longevity of placental MVNs for up to 14 days^9^. The magnitude of flow rate and its corresponding velocity and wall shear stress (WSS) are known to influence vascular diameters^12^. Static culture and low flow perfusion of MVNs results in regression and decreases in vascular diameters over 14 days^9^, while relatively high flow supplied by a microfluidic pump results in increases in vascular diameters over two days of continuous perfusion^11^. The magnitude of flow rate is an important MPS design feature to sustain long term (>14 days) perfusion of MVNs. Furthermore, the vast effects of flow on the morphology and function of MVNs necessitate detailed characterization of long-term perfusion of MVNs.

Here we present an MPS with 3D MVNs perfused with a microfluidic pump for 51 days and note several important phenomena. Continuous flow recovers perfusion of regressed vessels, and in perfusing initially healthy vessels, we observe long term (weeks) remodeling of the MVNs. Computational fluid dynamics (CFD) modeling of the MVNs provides insights into determining how individual vessel segments will remodel. Furthermore, long-term flow is shown to reduce inflammation of the MVNs. Finally, bulk transcriptomic analysis reveals governing pathways in the remodeling MVNs. The full characterization of this long-term, continuously perfused MPS provides insights into vascular responses to flow in engineered platforms and can be used to study long-term diseases and to incorporate aspects of hemodynamic processes previously not captured by MPS.

## Results

### 3.1 Physiological flow rescues perfusion in regressed vessels

Vascular networks grown under static conditions begin to regress after about one week in culture and are therefore of limited use for long-term experiments. To address this, we examined the ability of physiological flow to recover networks that were on the verge of losing. MVNs were grown in an existing device^11^ and flow was provided by a microfluidic pump^13^ to introduce flow at a specified time (**Fig. 1 a-c**). The pulsatility index, which is the difference between the maximum and minimum velocities divided by the average velocity, of the steady state flow was 0.30 (**Fig. 1 d**), comparable to the *in vivo* pulsatility index of cerebral mouse microcirculation^14^. Notably, the pump input pressure was held constant at 6 kPa for all flow experiments, which yielded the flow rates and pressure drops shown in **Supp. Fig. 1**. In brief, the flow rate was <10 μL/min and increased to ∼140 μL/min as the hydraulic resistance of the MVNs decreased, which corresponds roughly to an pressure difference of ∼1000 Pa and decreasing to ∼250 Pa. Network morphology, perfusability, and permeability were chosen as the three key features that could be compared directly to *in vivo* microcirculation.

**Figure 1.**
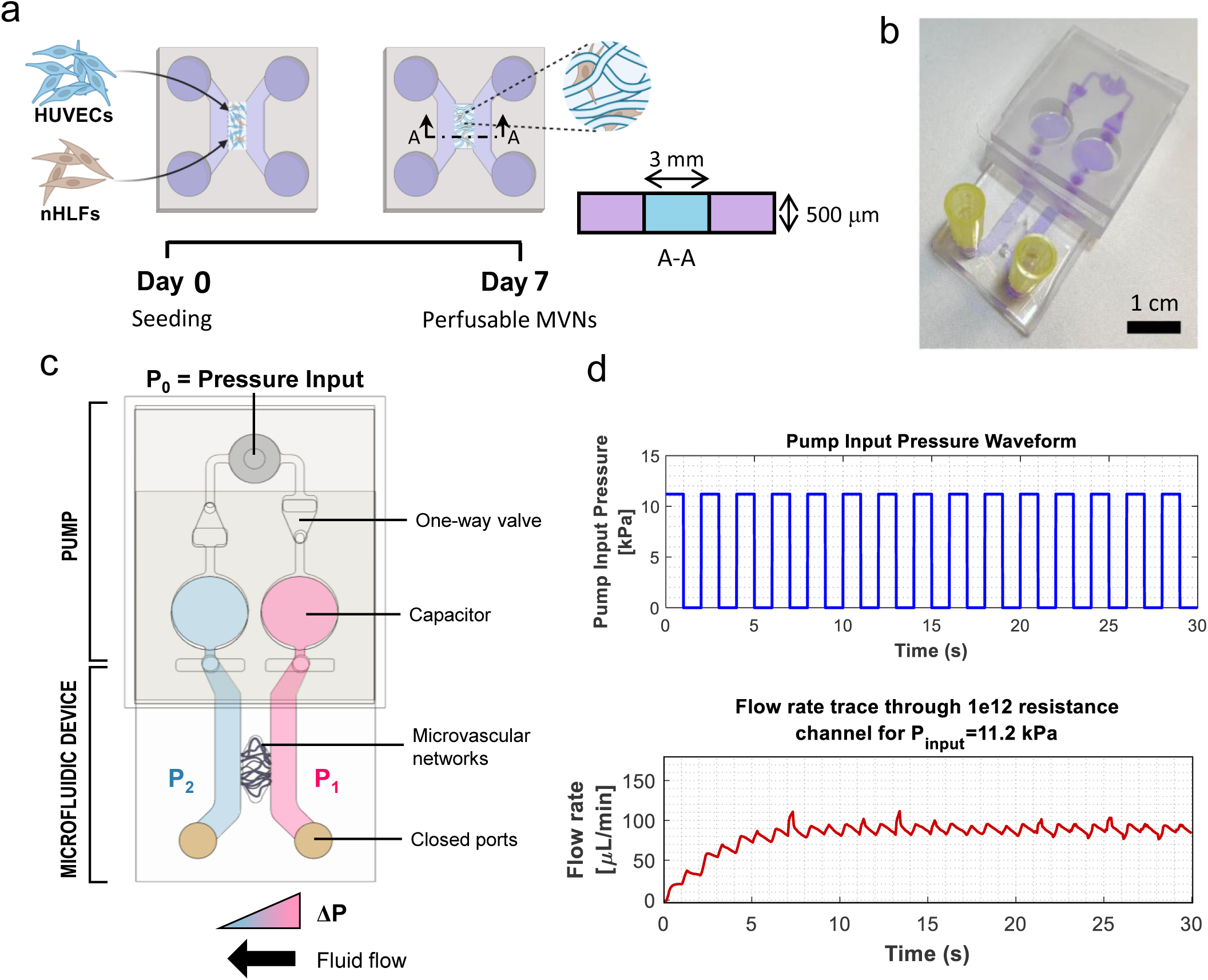
Experimental timeline and setup used for pump flow experiments. (a) The timeline for seeding and MVN formation in a defined microfluidic device. (b) A microfluidic pump is attached to the microfluidic device to introduce flow. (c) The pump generates recirculating flow by a pressure drop across the microvascular networks. Pump components are labelled. (d) The input pressure profile to the pump oscillates between a user-defined peak pressure and atmospheric pressure at a user-defined frequency. The corresponding bulk flow rate (trace is a moving average over 250 ms) quickly reaches steady flow within seconds of the input pressure commencing.

Visually, MVNs under static conditions were fully formed and interconnected on day 7 (**Fig. 2 a**) and then began to narrow, lose connections and, ultimately, cease to be perfusable (**Fig. 2 a**). Under static conditions, most MVNs were perfusable by day 7 but then regressed and lost perfusion by day 12 (**Fig. 2 a-c**). Flow commenced on day 12 and by day 15 the perfusable network fraction significantly increased compared to day 12 (**Fig. 2 a-c**). During the regression period (days 7-12), no differences in average MVN morphology were observed, likely due to measuring the average of the morphological features (**Fig.** 2 **d-g**). During the subsequent flow period (days 12-22), the average diameter (**Fig. 2 d**) and average length (**Fig. 2 e**) increased, the vascular density decreased (**Fig. 2 f**), and the vessel projected area (projected onto the x-y plane) increased until day 15 (**Fig.** 2 **g**). The permeability of the recovered MVNs on day 22 (10 days with flow) was comparable to MVNs on day 7 (**Fig. 2 h**). These results demonstrate that *in vitro* vascular networks can be rescued up to, and even beyond the time at which they lose perfusability. Thereafter, remodeling continues while vascular permeability remains relatively constant.

**Figure 2.**
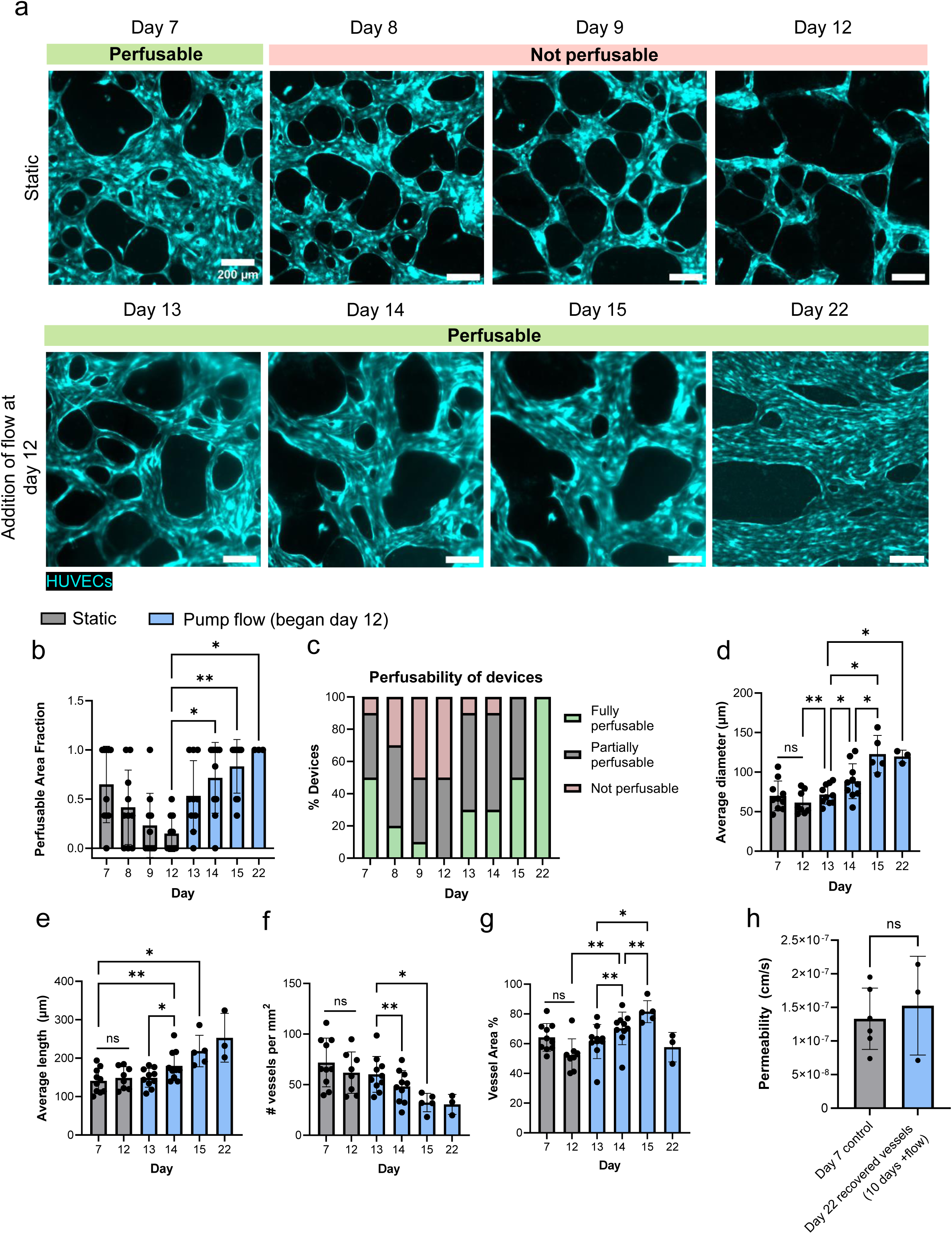
Flow can rescue perfusion in regressed vessels. (a) Representative images of the same vascular bed over time. Scale bar is 200 μm. (b) The perfusable area fraction of devices and (c) the collective perfusability of MVNs is reported. The average vessel diameter (d), length (e), vessel density (f), and vessel area (g) are reported. (h) The permeability of the recovered vessels on day 22 and control vessels on day 7 were measured. n = 3 – 10 MVNs.

### 3.2 Luminal flow increases the longevity of MVNs

Next, the question of how long the effects of flow can maintain barrier integrity was addressed. For all subsequent experiments, flow commenced on day 7. Applying flow to the MVNs increases their longevity but the MVNs lose resemblance to *in vivo* vasculature (**Fig. 3 a**). Media composition was found to play a role in the morphological remodeling of the MVNs in response to flow and growth factors, which can lead to uncontrolled growth (**Fig. 3 a**). To counteract this, medium without the supplemental growth factors found in growth medium, termed maintenance medium from here on, was used and the MVNs were perfusable for at least 51 days when experiments were stopped, though the system could have persisted for longer time (**Fig. 3 a**). Conversely, MVNs cultured with maintenance medium under static conditions regressed and lost perfusion by day 12 (**Supp. Fig. 2**), similar to culture with regular growth medium under static conditions. In view of these findings, all subsequent flow experiments used growth medium for the first 7 days during vessel growth and switched to maintenance medium from the time that flow was started on day 7 unless otherwise specified.

**Figure 3.**
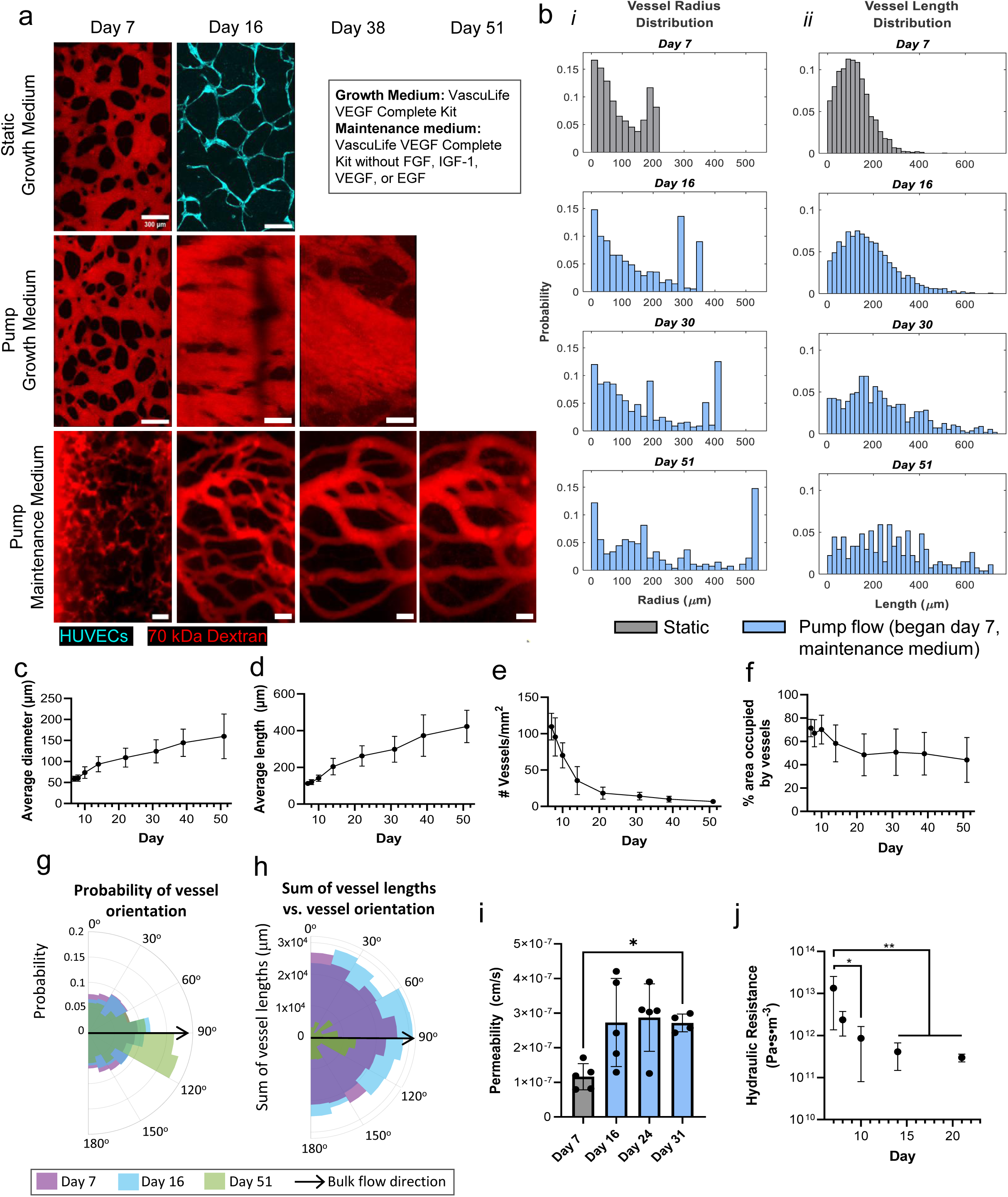
Pump flow increases the longevity of MVNs and vessels align with the direction of flow. (a) Representative images of MVNs grown statically, with flow and growth medium, or with flow and maintenance medium are shown perfused with dextran at selected timepoints. Scale bar is 300 μm. All the following figures are for MVNs with maintenance medium and pump flow (b-j). (b) The distribution of vessel radius (i) and length (ii) on days 7, 16, 30, and 51. The average diameter (c), average length (d), vascular density (e), and % area occupied by the vessels (f) are reported from day 7 to day 51. (g) Vessel orientation probability and (h) the orientation of the sum of vessel lengths is reported. (i) Permeability and (j) hydraulic resistance of the vessels over time. n = 4 – 11 MVNs.

Under these conditions, the MVNs continued to remodel over the course of 51 days. First, the distribution of the radii and lengths of the vessels was examined (**Fig. 3 b**). Prior to commencing flow, smaller vessel radii were more likely, and as the MVNs remodeled due to flow, the distribution of vessel radii increased to almost uniform on day 51, where large and small vessel radii were similarly likely to be present (**Fig. 3 b**). Notably, at all timepoints, there was a peak in the distributions at large radii; this was an artifact of the segmentation and skeletonization algorithm used that sets a limit on the maximum allowable vessel radius to avoid erroneous measurements. Similarly, the distribution of the vessel’s lengths followed a truncated Gaussian distribution with a bias towards shorter vessels prior to flow and then the distribution flattened and broadened until day 51, where long vessels were equally likely to be present as short vessels (**Fig.** 3 **b**). During the 44 days of flow, the average vessel diameter increased ∼3-fold (**Fig. 3 c**) and the average vessel length increased ∼4-fold (**Fig. 3 d**). The vascular density sharply declined in the first two weeks of flow, and by day 51 there was an 11-fold reduction compared to day 7 (**Fig. 3 e**). The area occupied by the vessels declined in the first week of flow and then remained steady (**Fig. 3 f**). Notably, vessel orientation changed dramatically over time with continuous flow (**Fig. 3 g**). From day 7 to 15, vessels had little directional orientation, but by day 51, most vessels became aligned with the direction of the pressure drop across the MVNs. The sum of the vessel lengths exhibited this same trend, but also demonstrated a large degree of vascular pruning with a dramatic reduction overall in vascular lengths (**Fig. 3 h**). For both probability and sum of vessel lengths cases, only small differences were noted between days 7 and 16, but by day 51 most of the MVN length was oriented in the direction of the pressure drop across the MVNs. These results indicate significant directional remodeling between days 16 and 51. Thus, while the rate of change in vascular morphology slowed over time, some changes continued even by day 51, suggesting that the MVNs remained dynamic.

The morphology of MVNs depends on the composition and structure of their surrounding matrix. The structure of a fibrinogen matrix depends on the concentration of fibrinogen^15^, therefore the of initial fibrin concentration on MVN morphology with flow was investigated. This was tested by seeding MVNs in either 3 mg/mL (control) or 5 mg/mL fibrinogen (high) (**Supp. Fig. 3**). On day 7, MVNs seeded in high fibrinogen appeared less connected and had smaller radii (**Supp. Fig. 3 a**) and were only partially perfusable (**Supp. Fig. 3 b**) compared to MVNs seeded in control fibrinogen. After 6 days of flow (day 13), all MVNs were fully perfusable, visually appeared similar (**Supp. Fig. 3 a**) and exhibited no significant differences in morphological parameters (**Supp. Fig. 3 c-f**), with the exception of vessel area coverage that was reduced by ∼ one third in high fibrinogen (**Supp. Fig. 3 c**). These results show that the perfusability of MVNs can be altered by initial seeding conditions such as fibrinogen concentration, but that continuous flow for six days minimizes these differences.

The permeability of the vessels remained low between days 7 and 31, ranging between 1 × 10^−7^ cm/s and 3 × 10^−7^ cm/s for 70 kDa FITC-dextran (**Fig. 3 i**), on the order of previously reported values^10^. The hydraulic resistance of the MVNs, which is the resistance to fluid flow, was ∼ 1 × 10^13^ Pa•s•m^−3^ on day 7 when the networks first became fully perfusable, then proceeded to decrease during one week of flow and remained stable between 1 × 10^11^ and 1 × 10^12^ Pa•s•m^−3^ until day 22 (**Fig. 3 j**). Luminal flow prolongs the life of MVNs, over which time the MVNs morphologically remodel while maintaining low values of permeability.

### 3.3 Unidirectional flow maintains vascular perfusability better than flow with alternated direction

Intravascular flow exerts a large effect on network morphology and function. The effect of flow with alternated direction was explored, as used in some previously reported experiments incorporating hydrostatic pressure to generate flows on a rocker platform would have the same effect (**Supp. Fig. 5 a**) ^16^. Three conditions were compared: 1) MVNs with rocker flow and with growth medium, 2) MVNs with rocker flow and with maintenance medium, and 3) MVNs with pump flow and maintenance medium. It is noteworthy that the two experiments also differed in terms of the peak or mean pressure drop across the MVN. For MVNs with a hydraulic resistance of 1e12 Pa•s•m^−3^, the peak pump flow rate was ∼70 μL/min while the peak rocker flow rate was ∼0.6 μL/min. Prior to flow, on day 7, all MVNs looked similar (**Supp. Fig. 5 b**). By day 16 (9 days of flow), the MVNs on the rocker with growth medium remained perfusable, but were narrow and did not appear to be aligned with the bidirectional pressure gradient (**Supp. Fig. 5 b**). Conversely, the MVNs on the rocker with maintenance medium regressed and lost perfusion by day 16 (**Supp. Fig. 5 b**). The MVNs with unidirectional flow and maintenance medium, however, were perfusable and appeared to align with the direction of flow by day 16 (**Supp. Fig. 5 b**). The stark difference between the perfusability and morphology of the MVNs between the rocker and pump flow using maintenance medium indicates that both the magnitude of flow and the directionality of the flow has significant effects on the long-term perfusability of MVNs.

### 3.4 Computational modeling reveals the role of flow rate, flow velocity and shear stress in the remodeling of MVNs

A computational fluid dynamics (CFD) model of the MVNs was used to analyze differences in flow characteristics between stable and regressing vessels. Analysis was performed on vessels at day 14 and vessel segments were categorized as either stable or regressing by observing whether they had regressed by day 21 (**Fig. 4 a**). The average radius of the stable vessels (150 +/- 69 μm) was significantly higher than that of regressing vessels (40 +/- 24 μm) (**Fig. 4 b**). There was no difference in the average wall shear stress (WSS) on stable or regressing vessels (**Fig. 4 c**). The average flow velocity (**Fig. 4 d**) and flow rate (**Fig. 4 e**) were significantly higher in the stable vessels compared to regressing vessels. Since vessel radius plays a significant role in the level of flow a vessel receives, the correlation between vessel radius and the various flow parameters was analyzed (**Fig. 4 f, g**). In the regressing vessels, there was a strong and significant correlation between vessel radius and velocity (r = 0.5693), and radius and flow rate (r = 0.8323) (**Fig. 4 f**). In the stable vessels, there was a significant and strong correlation (r = 0.6253) between the vessel radius and flow rate (**Fig. 4 g**).

**Figure 4.**
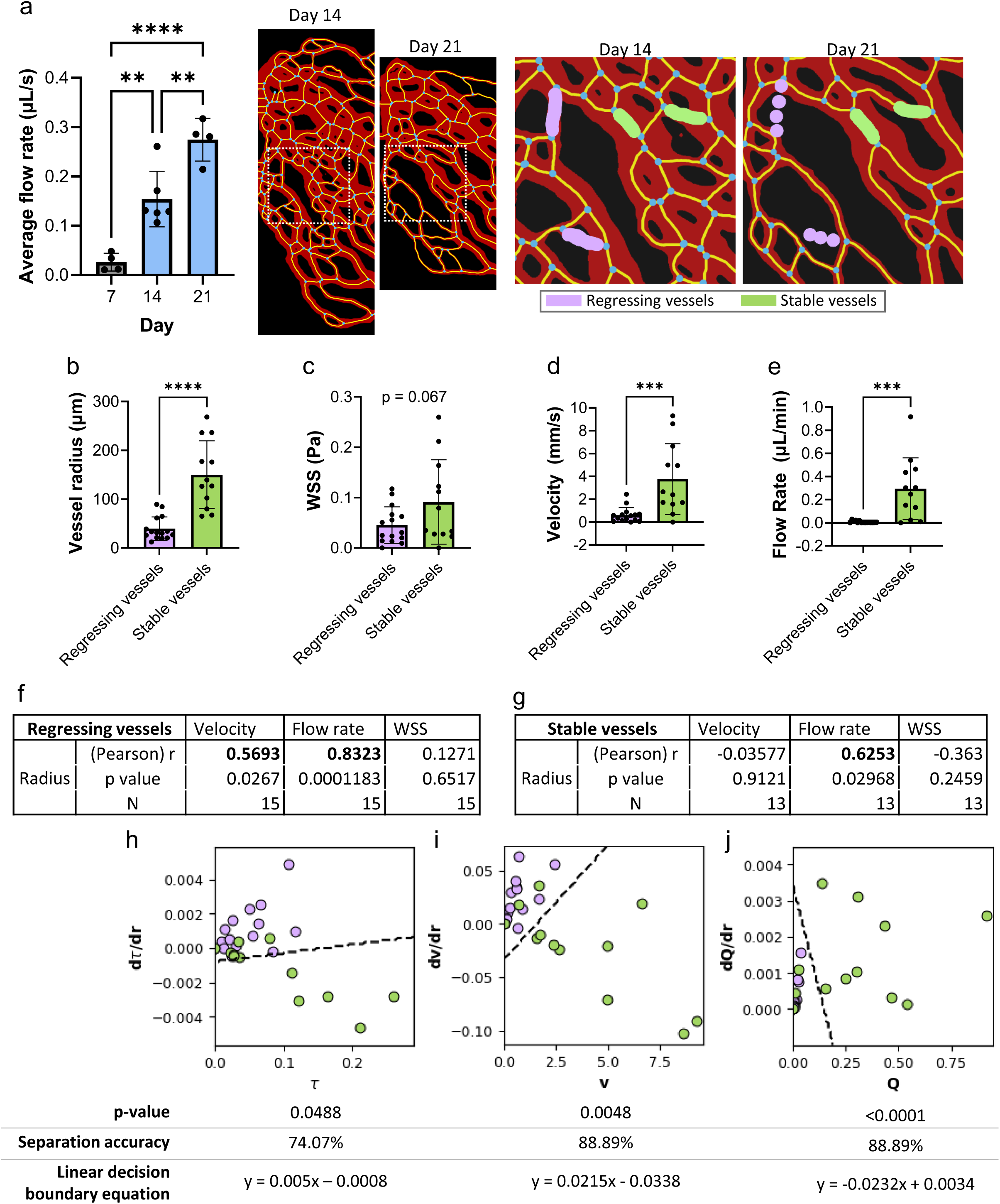
Stable vessels have different flow characteristics than regressing vessels. (a) Stable and regressing vessels were identified by comparing segmented images of vessels between day 14 and 21. Subsequent flow simulations and analysis were performed on the categorized day 14 vessels. The vessel radii (b), WSS (c), velocity (d), and flow rate (e) among stable and regressing vessels was compared. (f-g) A correlation analysis between vessel radius and velocity, flow rate, or WSS was conducted. A radius perturbation analysis was conducted and the change in WSS (h), velocity (i), or flow rate (j) with respect to change in radius vs. the original flow properties is reported, along with results from a 2D permutation test and linear decision boundary analysis. n = 12-15 vessel segments from 3 MVNs.

A CFD model was then used to systematically perturb the vessel diameters one at a time, decreasing their diameter by 10% and observing changes in WSS (τ), velocity (v), and flow rate (Q) (**Fig. 4 h-j**). Most regressing vessels experienced a decrease in their WSS (**Fig. 4 h**) and velocity (**Fig. 4 i**), while most stable vessels experienced an increase in their WSS (**Fig. 4 h**) and velocity (**Fig. 4 i**). All vessels experienced a decrease in flow rate because of the perturbation (**Fig. 4 j**). The vessels were compared in 2D space: one dimension was the change in flow parameter (ie. τ, v, Q) with respect to the change in radius, and the second dimension was the absolute value of the flow parameter. A 2D permutation test showed that the regressed vessels were significantly different than the stable vessels for all three flow parameter space comparisons (**Fig. 4 h-j**). The separation accuracies of a linear decision boundary analysis were about 74% for the τ space and about 89% for the v and Q spaces (**Fig. 4 h-j**). CFD modeling revealed that flow rate plays a critical role in the remodeling process of the MVNs.

### 3.5 Continuous perfusion reduces vessel inflammation

The supernatant from MVNs at days 7 (static), 14 (one week of flow), and 22 (two weeks of flow) was collected after 24-hours of conditioning with the MVNs and analyzed in a multiplexed inflammatory cytokine assay (**Fig. 5 a**). The concentration of GM-CSF (**Fig. 5 b**), IL-1β (**Fig. 5 c**), and IL-8 (**Fig. 5 g**) significantly decreased with 2 weeks of flow, indicating the MVNs became less inflamed. These cytokines are known to have various functions in the immune response: GM-CSF has been shown to act on macrophages and neutrophils by increasing their survival and activation^17^, IL-1β is known to induce expression of VCAM-1, which in turn contributes to leukocyte recruitment^18^, and IL-8 primarily acts to recruit monocytes and macrophages to an inflamed site^19^. GM-CSF, IL-1B, and IL-8 may be secreted from both ECs and fibroblasts (FBs) ^17,19,20^. While the other analyzed cytokines are known to play a role in vascular inflammation and remodeling, their concentrations did not decrease significantly with flow and time (**Fig. 5 d-f, h-k**). While the concentration of IL-6 tended to decrease over time, the differences did not reach statistical significance. This may be due to its role in chronic inflammation and tissue repair, whereas IL-8 is primarily associated with acute inflammation^21^. Furthermore, on day 7, growth medium and static conditions were used, while on days 14 and 22 maintenance medium and flow conditions were used, therefore the changes in cytokine concentrations were attributed to both the change in medium and the introduction of flow. These findings underscore the differential concentrations of inflammatory cytokines in response to continuous perfusion.

**Figure 5.**
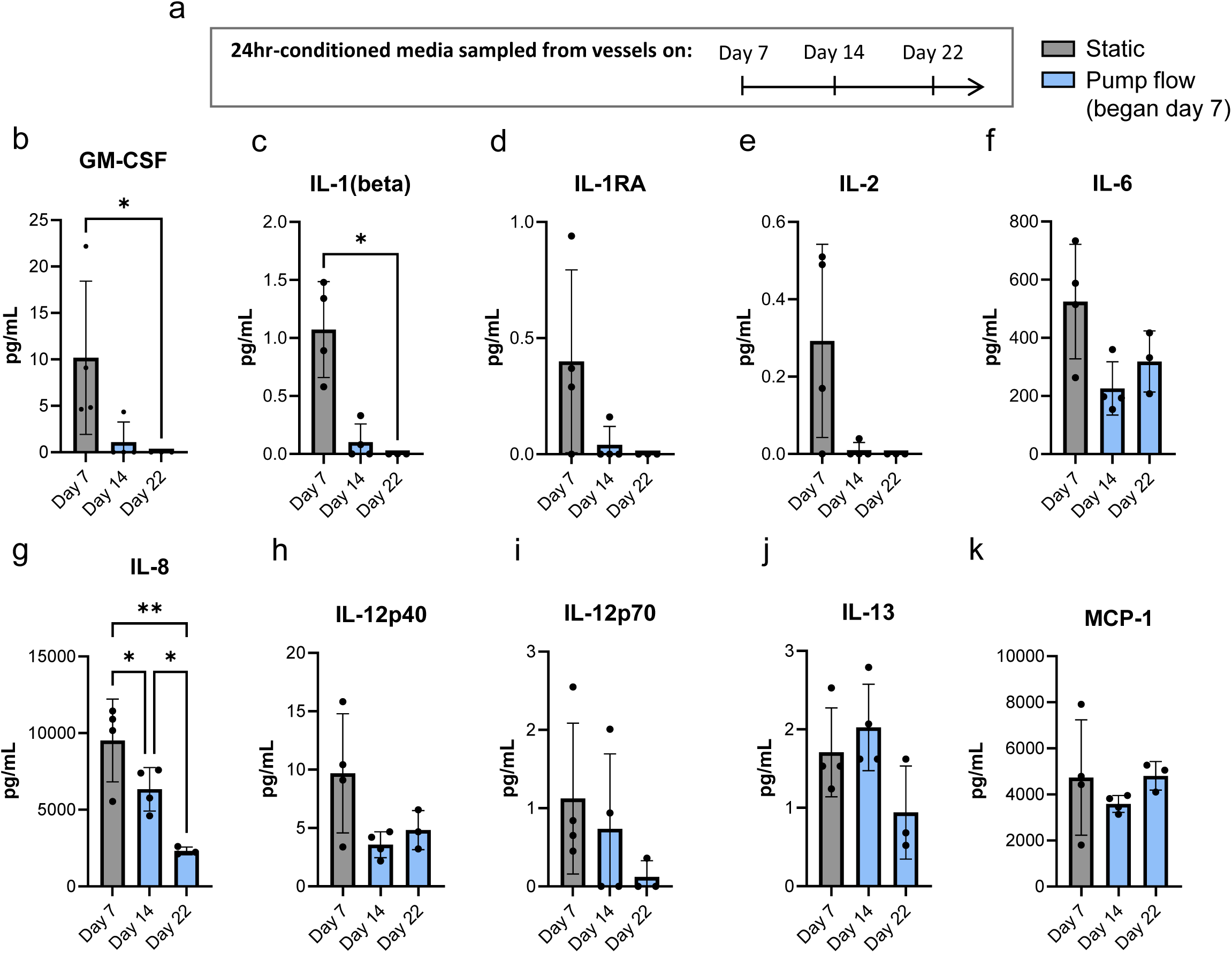
Flow reduces the magnitude and variability of the concentration of selected inflammatory cytokines. (a) Experimental timeline indicating the supernatant sampling timepoints. (b-k) Concentration of inflammatory cytokines across timepoints: days 7, 14 (7 days pump flow), and 22 (15 days pump flow). The selected cytokines were (b) GM-CSF, (c) IL-1β, (d) IL-1RA, (e) IL-2, (f) IL-6, (g) IL-8, (h) IL-12p40, (i) IL-12p70, (j) IL-13, and (k) MCP-1. n = 3-4 MVNs.

### 3.6 Bulk RNA sequencing provides an assessment of MVN response to flow

To identify differentially expressed genes (DEGs) as a function of duration of flow conditioning, the transcriptomes of the ECs and FBs from MVNs at days 7, 15 (8 days of flow), and 24 (17 days of flow) were analyzed using bulk RNA sequencing (RNAseq) (**Fig. 6 a**). The pairwise comparisons demonstrated distinct clustering patterns (**Fig. 6 b-c**). Volcano plots for the pairwise comparisons show the up-regulated and down-regulated (red) DEGs (p.adjust < 0.05, |log_2_ fold change| > 1) (**Fig. 6 d**). *ICAM-1* was downregulated in ECs at day 24 compared to days 15 and 7, and *VCAM-1* was downregulated in ECs at day 15 compared to day 7, potentially indicating a less inflamed state. Inflammation triggers increased expression of ICAM-1 and VCAM-1, which facilitate leukocyte transendothelial migration ^22–24^. *VCAM-1* gene expression has been shown to be inhibited in regions of laminar flow, mediated through shear stress-regulated genes such as endothelial nitric oxide synthase or superoxide dismutase^18^. mRNA expression of *HMOX1*, *GCLM*, and *NQO1* was upregulated in ECs at day 15 and day 24. HMOX1, GCLM, and NQO1 proteins have been shown to be upregulated in ECs exposed to atheroprotective shear flow ^25,26^. The total number of DEGs is reported (**Fig. 6 e**). The day 24 vs. day 7 pairwise comparisons yielded the highest number of DEGs in both the ECs and the FBs. Venn diagrams show the grouped overlap of up- and down-regulated DEGs among the three pairwise comparisons for ECs and FBs (**Fig. 6 f**). Interestingly, there were no DEGs only common between the day 24 vs. day 15 and day 24 vs. day 7 pairwise comparisons for the ECs or the FBs, indicating continuous transcriptomic changes in the cells. 8 DEGs were common to all three EC pairwise comparisons, and 13 DEGs were common to all FB pairwise comparisons. *ITGB5*, common among all EC pairwise comparisons, encodes integrin subunit beta 5, has been shown to play a role in lung endothelial survival and migration^27^. *SRXN1* was common among all EC and FB pairwise comparisons and its encoded protein has been shown to protect neurons^28^ and cardiac progenitor cells^29^ from oxidative stress injury. GO pathway analysis (**Fig. 6 g**) revealed differential expression of hypoxic pathways in both the ECs and FBs, which may point to the differential expression of *SRXN1* in the ECs and FBs. *EPHX1*, which has been shown to have detoxification properties^30^, was also upregulated in all pairwise comparisons. Among the FB pairwise comparisons, *GPX3* and *ADM* were common DEGs of interest. *GPX3* is known to be upregulated in oxidative stress conditions and protects epithelial cells from oxidative damage^31^. Adrenomedullin, the protein encoded by *ADM,* has been shown to induce vasodilation^32^, which is in line with the MVN morphological changes presented here. The GO pathway analysis further revealed that by two weeks of flow conditioning (day 24 pairwise comparisons), the ECs were involved in pathways related to cell adhesion, lung development and ECM organization, specifically collagen fibril organization (**Fig. 6 g**). Several collagen genes were upregulated by the ECs, namely *COL1A1*, *COL1A2*, and *COL12A1.* The FBs were also involved in extracellular matrix reorganization, collagen fibril organization, and blood vessel development (**Fig. 6 g**). By two weeks of flow conditioning, several hypoxia-related pathways were differentially expressed in the FBs, indicating a possible lack of oxygen (**Fig. 6 g**). Perivascular FBs in a zebrafish mode have been shown to help regulate blood vessel diameter through the secretion of collagens^33^. The “collagen fibril organization” GO pathway was differentially expressed and several collagen-related genes were significantly upregulated in the FBs at d24vsd7, including *COL11A1*, *COL27A1*, *COL5A2*, and *COL14A1*, which may indicate increased collagen production by the FBs. Furthermore, several pathways related to biosynthetic and metabolic process were differentially expressed in the FBs at d24vsd7, which may further indicate excessive collagen production by the FBs. FB production of collagen is known to be energy intensive and requires unique biosynthetic demands of the FBs^34^. The RNAseq results provide insights into the dynamics of the EC and FB phenotypes over long term MVN perfusion.

**Figure 6.**
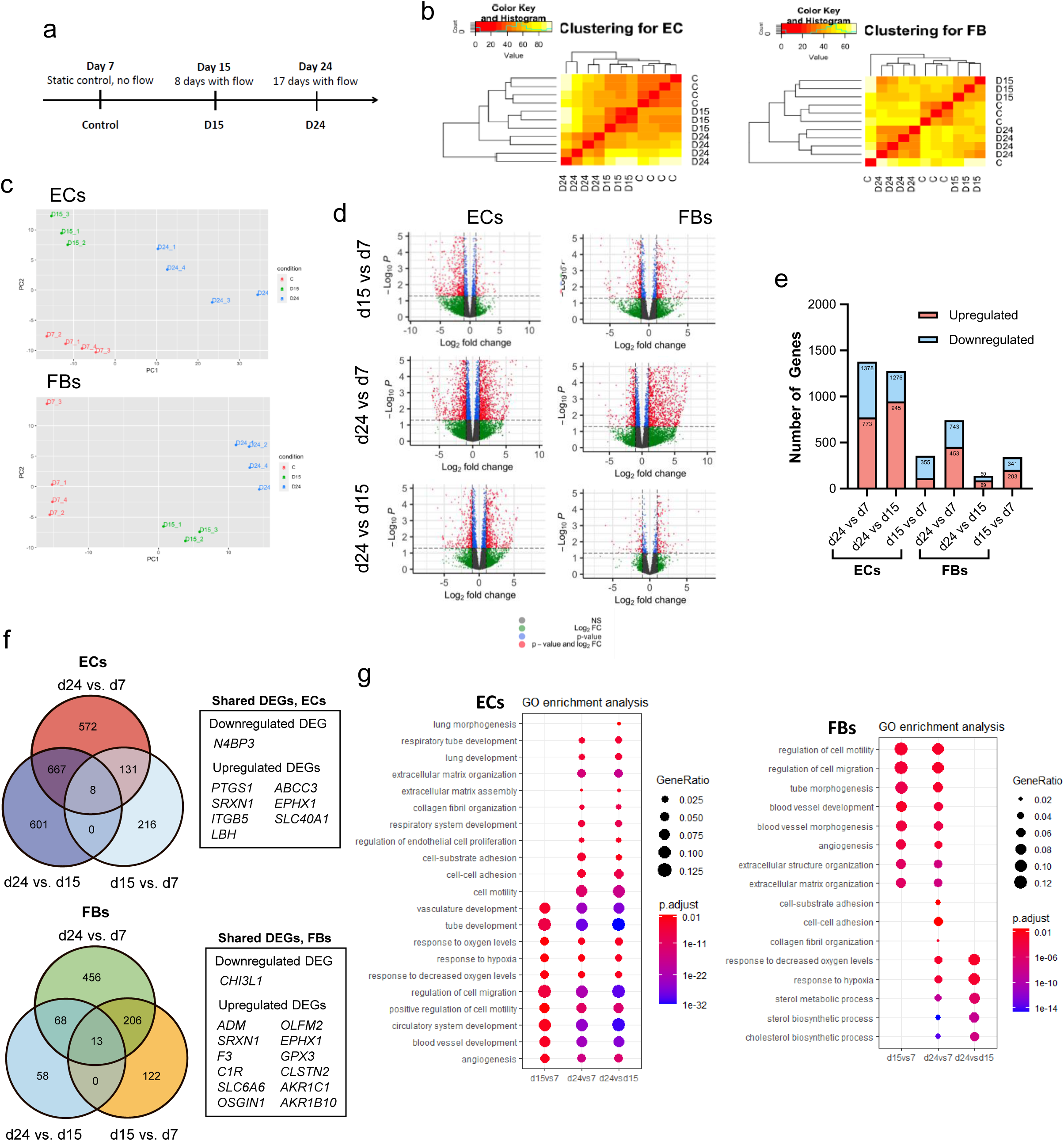
Bulk RNAseq of the ECs and FBs revealed differential pathway expression between days 7, 15 (8 days pump flow), and 24 (17 days pump flow). (a) Cells were isolated and prepared for RNA extraction on days 7, 15, and 24. (b-c) Heatmaps (b) and PCA plots (c) showing clustering of the ECs and FBs for each pairwise comparison. (d) Volcano plots for genes in the ECs and FBs for each pairwise comparison with annotated DEGs of interest. (e) Number of DEGs up- and down-regulated in the ECs and FBs. (f) Venn diagrams showing the overlap of DEGs among the pairwise comparisons. Inset table lists the DEGs common to all three pairwise comparisons for the ECs and FBs. (g) GO pathway analysis comparing the ECs and FBs across the pairwise comparisons timepoints. n = 3-4 samples pooled with 3-4 MVNs.

## Discussion

The purpose of this work was to investigate the effects of continuous recirculating flow through MVNs grown in microfluidic devices. These systems have been used for studies of primary and metastatic cancer^35^ and for various organ-on-chip models ^36^. However, these networks have a limited lifetime, and the useful period of experimentation is typically several days after the MVN forms. But in many applications, such as studies of metastatic outgrowth or organ maturation and function, longer-term studies are required. In the present study, flow is used to recover the perfusability of regressing MVNs, which subsequently with continuous flow maintain perfusion for at least 51 days (**Fig. 7**). This enabled us to observe how MVNs exposed to physiological flow remodel through a combination of vessel dilation and regression and how their phenotype and state of inflammation change under flow compared to standard, static conditions.

**Figure 7.**
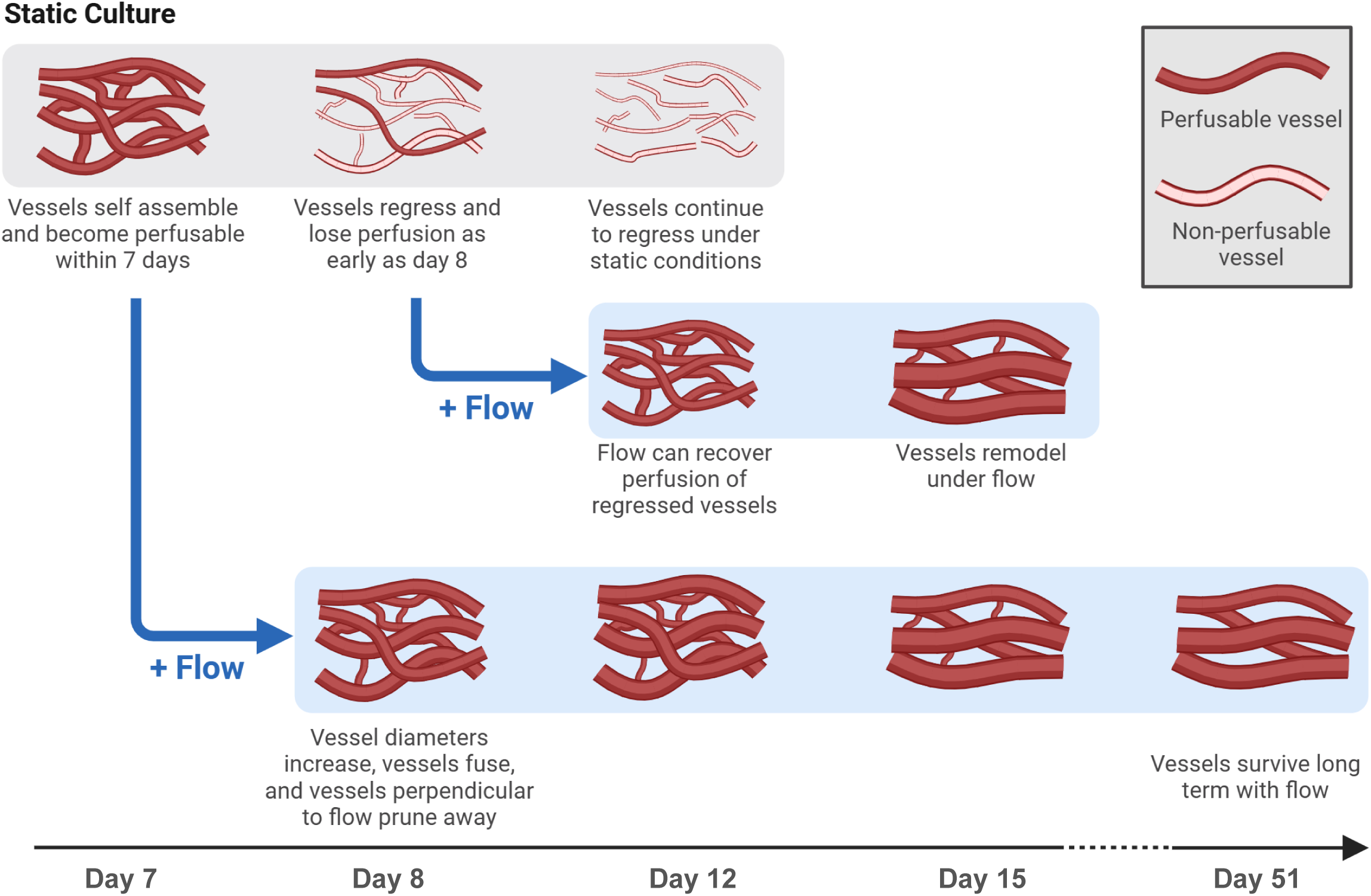
Summary of the effects of flow on the remodeling of MVNs. Under static culture, MVNs lose perfusion, regress, and die. Adding flow to regressing vessels rescues perfusion and vessels remodel in response to flow. Adding flow to initially healthy and perfusable vessels results in vascular remodeling and long-term perfusion (for at least 51 days).

Over the course of 44 days of flow, regression of vessels perpendicular to the pressure gradient was observed, consistent with previous findings ^37^, whereas the diameters of vessels parallel to the pressure gradient increased over time. The permeability of the MVNs remained low. A previous study using a similar MPS reported similar permeability values for MVNs under hydrostatic pressure-driven flow^10^. The higher permeability on Day 31 may have been a result of a lower effective dextran concentration^38,39^ because of the method used to introduce dextran when the pump was connected to the device. While significant remodeling was anticipated, it was also expected that the networks would eventually stabilize based on a long history of observations *in vivo*. This expectation is the basis for what has come to be known as the set point theory^40^, wherein vessels experiencing higher than physiological WSS tend to increase in diameter until the WSS attains a certain value. Conversely, vessels with a WSS less than the set-point value constrict. This is but one of many biological consequences of endothelial shear stress, which has been studied extensively both *in vivo* and *in vitro* in the context of arterial disease (see ^12^ for a comprehensive review). For example, in response to an arteriovenous shunt in canine observed 6-8 months post-operatively, the radius of the shunt vessel increased with increased flow load until the wall shear rate reached the control, pre-shunt value^41^. This may suggest that longer culture time is required for the MVNs to reach stability. Furthermore, it has been shown in a zebrafish model that peak WSS imposed by individual red blood cells through capillaries can provide sufficient mechanical stimulus for microvascular vessels to reach their stable radius^42^. The MVNs in this study, however, did not approach an average set point.

There are several possible explanations why the average diameters of the MVNs continued to increase. The continued expansion of vessel diameter observed may have been due to the presence of a high concentration of growth factors in the medium as the progressive dilation was significantly reduced when using maintenance medium. When the growth factors were removed, major vessel segments tended to stabilize maintaining a nearly constant diameter between days 16 and 51 and vascular network resistance remained nearly constant on days 14 and 22. Changes in stiffness and composition of the surrounding gel may also have had an effect on the changing vascular morphology^43^. In a static 3D *in vitro* platform similar to the one presented in this study, the elastic modulus of the hydrogel with embedded ECs and FBs increased approximately 13-fold in the first seven days of juxtacrine co-culture and thereafter maintained a somewhat constant elastic modulus through day 14^4^. This same static system was used for proteomics analysis by mass spectrometry, which showed that within 14 days of seeding, 33% of the matrix was newly deposited by the cells (Whisler, J. Engineered, functional, human microvasculature in a perfusable fluidic device. *DSpace@MIT* https://dspace.mit.edu/handle/1721.1/113761 (2017)). The newly synthesized matrix was comprised of a wide variety of matrix proteins especially several types of collagens (COL1A1, COL1A2, COL6A1 COL6A2, COL6A3, COL12A1). Interestingly, the large changes in matrix synthesis were not observed when the endothelial cells or fibroblasts were individually cultured under the same conditions, suggesting that bi-directional signaling between the two cell types is critical for these changes. A similar trend would be expected for the flow system presented in the present study as the same two cell types were used. Other matrix proteins that were clearly present in the static experiments, however, notably fibrinogen (FN1) and heparin sulfate (HSPG2), were only slightly up- or down-regulated, respectively, under flow conditions. The lack of pericytes and smooth muscle cells may also contribute to the continued increase in diameter. Pericytes are known to regulate vascular diameter and vascular blood flow in the capillary bed through vasoconstriction and vasodilation ^44^, and smooth muscle cells are known to play a role in vasoconstriction and vasorelaxation in the arterioles^45^. The heterogeneity of the MVNs may have contributed to the continuous vascular remodeling, whereby the initial morphology and structure of the MVNs at day 7 was not the optimal organization for continuous flow perfusion. The transcriptomic data suggest yet another explanation. Between days 7 and 24, there is a strong indication that the fibroblasts are hypoxic, even under continuous flow. If so, the physiological response would be to signal an increase in perfusion, which would correspond to an increase in vessel diameter. Vasodilation and constriction is a multifaceted, dynamic process, that depends on many factors in addition to WSS.

MVNs perfused with flow with alternated direction, conversely, did not show increases in vessel diameter over time. Despite the extensive literature on the effects of pulsatile or flow with alternated direction on endothelial function in the context of arterial disease, relatively little research has addressed these effects in the microcirculation. This is likely due, at least in part, to the fact that flow with alternated direction is uncommon in the small vessels under normal physiological conditions since flow in the capillaries is generally thought to be unidirectional and, while pulsatile, the temporal fluctuations are small relative to those in the larger arteries. Despite this, rocker platforms are becoming increasingly common as MVN culture agents, and the effect of flow with alternated direction on MVNs must be characterized. The observed deterioration of the MVNs under rocker flow compared to continuous flow suggests that either the directionality of the flow is significant or that some minimum flow rate must exist for MVNs to maintain perfusability. Due to physical constraints, the magnitude of the pressure gradients generated by the rocker was typically lower than that in unidirectional flow (∼ 3.9×10^5^ Pa/m for the pump and ∼ 3.3×10^4^ Pa/m for the rocker) and this could well have contributed to the differences observed. This is further supported by the CFD simulations conducted on stable and regressing vessels, which showed that stable vessels had significantly higher flow rates than regressing vessels. As for physical mechanisms, one potential mechano- sensor would be the primary cilium present in most mammalian cells ^46^ including endothelial cells. This singular structure is known to elicit a variety of cellular signals, and has been shown to be responsible for preventing vessel regression, and could be capable of sensing the direction of flow^47^. This comparison highlights the significance of the magnitude and directionality of flow on the dynamics of MVNs.

Focusing now on stable vessels and regressing vessels rather than the average vessel, CFD modeling of flow through MVNs revealed several interesting findings. One study thus far has demonstrated long term perfusion for up to 50 days and showed that vessels located in regions of low flow were more likely to regress, but did not characterize the function or phenotypes of the vasculature ^37^. In addition to the apparent role of flow rate in determining the stability of a vessel, the tendency for regressing vessels to experience a decrease in their WSS when their diameters are reduced by 10% may indicate that their morphology and location relative to other vessels may also influence their stability. The application of either a constant flow rate or constant pressure drop across the MVNs may also influence their stability; e.g., an increase in vessel diameter under a constant flow rate decreases WSS, while under a constant pressure gradient the WSS increases^42^. However, the microfluidic pump used in this study is neither a constant pressure nor a constant flow rate pump. CFD modeling provided insights into morphological remodeling of the MVNs in response to flow.

In addition to influencing MVN morphology, flow also played a role in lowering inflammation of the MVNs. The cytokine concentrations on day 7 were sampled from MVNs before exposure to flow, which indicates that the initial state of the system is relatively inflamed. Initially higher concentrations of cytokines at day 7 may be a result of the use of a fibrin-based hydrogel used in cell seeding. Fibrin is known to be involved in inflammation and cytokine/chemokine production. IL-6 and MCP-1 were shown to accumulate in a fibrinogen-dependent manner in mice^48^. Data obtained in static conditions show that there is significant remodeling of the ECM over 14 days, which may decrease the presence of fibrinogen and thus reduce inflammatory effects. The concentration of several inflammatory cytokines subsequently decreased with 7 and 15 days of flow in maintenance medium, indicative of a gradual change toward a healthy state. This has several important consequences for the application of *in vitro* MVNs to model disease processes. The reduction in inflammatory cytokines is consistent with an atheroprotective flow, which is characterized by high unidirectional laminar shear stress^49^. Looking at specific downregulated cytokines, IL-8 significantly decreased over time. It is important in inflammation, whereby it enables adhesion of monocytes ^50^ and activation of neutrophils ^51^, so reduced levels of IL-8 can thus be associated with a less inflamed state. In a study using a static co-culture of HUVECs and normal human lung FBs in a fibrin gel in a microfluidic chip, the concentration of secreted IL-6 decreased from day 3 to 5, but there were no changes in IL-8 or MCP-1 ^52^, which may indicate that flow or longer culture time is required for secreted IL-8 to decrease. The reduction in IL-8, IL-1β, and GM-CSF also suggests a shift towards a less angiogenic state, consistent with our gene expression analysis. IL-8 is increased in the early stages of angiogenesis ^53^, and GM-CSF ^54^ and IL-1β ^55^ promote angiogenesis. This result is further supported by our findings showing a reduction in the gene expression in the ECs of *ICAM-1* between day 24 and days 15 and 7, and *VCAM-1* between days 15 and 7. IL-1β is known to induce VCAM-1 expression^18^. A reduction in IL-1β may contribute to the lower expression of *VCAM-1* seen here. Furthermore, MMP-2 has been shown to degrade IL-1β ^56^, thus the reduction in IL-1β in this system may be attributed to the increase in *MMP-2* mRNA expression in the FBs at day 24. High, sustained levels of MCP-1 are consistent with increased mRNA expression in the FBs at day 15 and day 24 compared to day 7 control. Overall, the system becomes less inflamed with flow over time, with some of the changes being attributed to the changing transcriptomics of the EC and FBs.

Transcriptomic analysis also revealed several differentially expressed pathways throughout the remodeling of the MVNs. The RNAseq analysis indicates that the MVNs are likely continuously remodeling throughout the 24 days, consistent with morphological and functional data presented earlier. The HUVECs are known to be plastic^57^, and may have been becoming more lung-like due to the long-term co-culture with the lung FBs. The continued differential expression of angiogenic and ECM pathways in the ECs may indicate that the ECs were continuously in a state of vascular remodeling. ECM remodeling-related pathways were differentially expressed in both the ECs and the FBs. In an MPS similar to the one presented here, 33% of the matrix was newly deposited by the ECs and FBs under static culture within 14 days of seeding (Whisler, J. Engineered, functional, human microvasculature in a perfusable fluidic device. *DSpace@MIT* https://dspace.mit.edu/handle/1721.1/113761 (2017)). Several genes encoding the proteins from the 33% newly deposited matrix were upregulated in the MVNs of this study, namely *COL1A1*, *COL1A2*, and *COL12A1in* the ECs. Both the ECs and FBs had several highly upregulated collagen-related genes, potentially indicating deposition of collagen in the remodeling ECM. Importantly, both the ECs and FBs contribute to ECM remodeling. Hypoxia-related pathways are differentially expressed in the ECs and the FBs. The ECs may be expressing hypoxic pathways because of the flow rate-mediated vascular remodeling, where the population of ECs lining regressing vessels that are receiving inadequate levels of flow rate may be hypoxic. The FBs may be expressing hypoxic pathways due to the reduction in vascular density from 35 vessels/mm^2^ at day 14 to 18 vessels/mm^2^ at day 22 (**Fig. 3 e**), which may be below some threshold for adequate oxygen supply. Cells in organs and tissues are typically within 200 μm from a blood vessel to have adequate nutrient and oxygen supply ^58^. The ECM of flow conditioned MVNs has lower diffusivity to 70 kDa Dextran compared to static MVNs, which may point to a reduction in oxygen diffusivity^9^. Further studies are required to identify methods to ensure the cells in the ECM are receiving adequate amounts of oxygen. Transcriptomic analysis revealed key pathways present in the remodeling MVNs in response to flow and pointed towards future directions of study.

While the findings provide valuable insights, the following limitations should be considered. The CFD model used to simulate flow through the MVNs has several inherent assumptions that may result in errors of the absolute flow estimates made here. First, the model assumes steady, laminar flow. The average Reynolds number (Re) calculated was found to be 2.6 and the maximum Re was 11.6, validating the assumption of laminar flow, but suggesting that some small effects of inertia could be present. More importantly, errors in the measurement of vessel diameters have a particularly strong effect on the estimated hydraulic resistance of the vessel segments owing to the inverse fourth power dependence of vessel flow resistance (*R*) on radius (*r*); *R = 8μL/πr*^4^. For example, vessel diameters were measured from maximum projections, 2D images on the x-y plane, which do not consider the two-to-one ellipticity ratio of the vessel cross sections (**Supp. Fig. 7**). Finally, flow sensitive genes such as KLF-2 did not show differential gene expression in the flow MVNs compared to the day 7 static controls. The half-life of KLF2 mRNA was previously reported to be 56 minutes ^59^, significantly shorter than the hours-long time required to digest, extract, and sort the cells from the microfluidic devices. Furthermore, the mRNA from the ECs was extracted after a relatively long flow exposure time, whereas in the literature ECs are typically exposed to shear flow for 16-72 hours ^60–62^.

This MPS extends upon existing models by enabling the capability to study hemodynamically modulated processes, such as vascular regression/reperfusion, drug distribution, or metastatic dissemination. Subsequent investigations might extend this research by incorporating organ-specific cells or patient-derived cells to study the role of hemodynamics in organ- and patient-specific diseases.

## Conclusion

Here, we use an inexpensive, low dead volume microfluidic pump to provide long-term, recirculating flow through MVNs. We show that continuous perfusion of a microvascular microfluidic system is capable of long-term (at least 51 days) perfusion. During perfusion, the MVNs remodel: thin vessels prune and vessels that align with the direction of flow predominate, and their diameter increases. The addition of flow to regressed MVNs is observed to recover perfusability to initially non-perfusable networks. Computational modeling suggests flow rate is a determining factor in the regression or stability of individual vessel segments. MVNs become less inflamed with continuous perfusion and are active in extracellular matrix reorganization, hypoxia, and vascular development transcriptomic pathways. These findings pave the way to developing long-term MPS and models to study complex hemodynamic processes such as drug distribution.

## Methods

### Cell culture

Immortalized human umbilical vein endothelial cells (ECs) (Lonza, CC: 2935, immortalized and transfected to express blue fluorescent protein as previously described ^63^), and human primary normal lung fibroblasts (FBs) (Lifeline, FC-0049) were used in this study. ECs were cultured with vascular growth medium using the vendor’s protocol (VascuLife VEGF Endothelium Medium Complete Kit, Lifeline) and used at passage 8; FBs were cultured with fibroblast medium following vendor’s protocol (FibroLife S2 Fibroblast Medium Complete Kit, Lifeline) and used at passage 5.

### Microvascular network formation

ECs and FBs were detached using Accutase (Sigma, SCR005), pelleted, and resuspended in vascular growth medium supplemented with 4 U mL^−1^ thrombin (Sigma, T4648-1KU) at concentrations of 26 × 10^6^ mL^−1^ and 6 × 10^6^ mL^−1^, respectively. Equal volumes of resuspended HUVECs and FBs were mixed, then combined with an equal volume of fibrinogen (6 mg mL-1, Sigma 341578) in phosphate buffered solution (PBS, Gibco 10010031), and injected into the gel ports of the microfluidic devices. The devices were placed in a humidified incubator for 12 minutes to allow the hydrogel to polymerize, and then vascular growth medium was added into the device medium channels. Medium in these channels was changed daily and MVNs became perfusable by day 7.

### Fabrication of microfluidic device and pump

The design of the microfluidic device and pump were created using Autodesk Fusion 360. The device was composed of a central gel channel flanked by two media channels. The gel channel was 3 mm wide, 7 mm long, and 0.5 mm tall, and the media channels were 3 mm wide and 0.5 mm tall. The pump design was used as described previously ^13^. The device and pump molds were milled using a Bantam Tools Desktop CNC Milling Machine (Bantam Tools). Polydimethylsiloxane (PDMS, Dow Corning Sylgard 184, Ellsworth Adhesives) was mixed at a 10:1 elastomer to curing mass ratio, degassed for 40 minutes, poured into the device and pump molds, degassed a second time for 25 minutes, then placed and cured at 65°C overnight. Individual devices and pumps were cut out, and the ports of the gel channel, media channel, and pump were punched using biopsy punches (Integra Miltex). The silicone membrane (LMS, Amazon) was cut to size and ports were punched using biopsy punches (Integra Miltex). The devices, pumps, and silicone membranes were sterilized in an autoclave for 25 minutes. The devices and #1 glass cover slips (VWR) were exposed to plasma (Harrick Plasma), bonded together, and placed in an 75°C oven overnight. The pumps were similarly bonded through plasma exposure in a two-step method: first the bottom half of the pump was bonded to the silicone membrane, and then the bottom half and silicone membrane group was bonded to the top half of the pump, and placed in a 75°C over overnight, as previously described ^13^.

### Application of flow to MVNs

The pump was connected to the device using silicone tubing (Miniature Firm EVA Tubing for Air and Water, 1883T3, McMaster Carr) on either day 7 (long-term experiments) or day 12 (perfusion recovery experiments). All flow experiments used a pump input pressure of 6 kPa. Long-term flow experiments were performed with maintenance medium after day 7 (VascuLife VEGF Endothelial Medium Complete Kit without rh VEGF LifeFactor, rh FGF basic LifeFactor, rh IGF-1 LifeFactor, and rh EGF LifeFactor, Lifeline), and perfusion recovery experiments were performed with growth medium (VascuLife VEGF Endothelium Medium Complete Kit, Lifeline) after day 12. During pump operation, the two device reservoirs opposite the pump were blocked to create a closed fluidic system. Media in pump devices was changed every two days.

The corresponding flow rate through and pressure drop across the MVNs was dependent on the hydraulic resistance of the MVNs and followed relationships according to **Supp. Fig. 1**. This input pressure was chosen based on computational simulations that predicted physiological average wall shear stresses of 1 Pa, as previously described ^11,13^. The steady state flow rate depended both on the pressure difference across the pump capacitors and the hydraulic resistance of the MVNs and channels connected to the pump. The steady state flow rate for different pressure inputs and characteristic hydraulic resistances was measured (**Supp. Fig. 1**). The hydraulic resistance was measured using a previously discussed protocol^11^. The corresponding pressure drop was calculated using

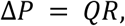

where Δ*P* is the pressure drop, *Q* is the steady state flow rate and *R* is the hydraulic resistance of the connected channels. The pump flow rate increased linearly with input pressure (**Supp. Fig. 1**).

A custom-built rocker was used to generate hydrostatic pressure-driven flow. 5 mL syringes were inserted into the microfluidic device ports. The syringes were all filled equally to achieve a height of 20 mm. The devices with syringes were then placed inside of a box with a dish containing PBS and placed on the rocker platform. The rocker tilted 43° such that one media channel was elevated higher than the second media channel, producing a hydrostatic pressure difference to drive flow through the MVNs (**Supp. Fig. 5**). This tilt was maintained for 10 minutes, after which point the rocker tilted 86° in the opposite direction to reverse the hydrostatic pressure difference and continued tilting or “rocking” every 10 minutes. Every 48 hours, the medium in the syringes was replaced with fresh medium. This setup produced a maximum hydrostatic pressure difference of 98 Pa, which dissipated to 96 Pa during the 10 minutes for MVNs with 1e13 Pa-s/m^3^ hydraulic resistance. This translates to an initial bulk flow rate of 0.59 μL/min and dissipated to 0.58 μL/min after 10 minutes. The hydrostatic pressure difference created by the rocker can be estimated following a similar analysis shown in **Supp. Fig. 6**, and, using the hydraulic resistance of the MVNs, the flow rate can be calculated as specified above.

The microfluidic pump features a low dead volume (230 μL), recirculating flow, and flow rates ranging from 8 – 240 μL min^−1^, allowing for continuous, long-term perfusion of microvascular networks with or without circulating cells at physiological levels of flow. The low dead volume allows for adequate buildup of secreted factors, such as cytokines.

### Imaging of MVNs, average morphological analysis, and permeability assay

MVNs were imaged using either an Olympus FV1000 (Olympus, Japan) confocal laser scanning microscope or a Nikon Eclipse Ti epifluorescence microscope (Nikon, Japan). Epifluorescent images of the MVNs were acquired using the 4x objective. Morphological analysis was performed in Image J (NIH) by thresholding the MVN signal using Trainable Weka Segmentation, measuring the area of the MVNs, skeletonizing the MVN, and analyzing the skeleton using “Analyze Skeleton”. Average MVN diameter was then calculated as

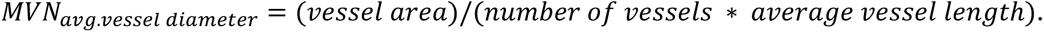

Permeability of MVNs under static conditions was measured by aspirating medium from the device media channels and introducing 100 μL of 0.1 mg mL^−1^ 70 kDa Texas Red dextran (Thermo Fischer Scientific). For flow-conditioned samples connected to the pump, 100 μL of 0.1 mg mL^−1^ 70 kDa Texas Red dextran (Thermo Fischer Scientific) was added to the downstream media reservoir, the pump was manually actuated twice to flow in the dextran, and the excess fluid from the upstream reservoir was aspirated. Then, the change in intravascular and extravascular fluorescence intensity was quantified, as previously described ^64^. Confocal z-stacks were acquired every 12 minutes using the confocal microscope with the 10x objective at 640 px resolution and 5 μm steps in the z-direction. The change in fluorescence intensity was analyzed using an ImageJ (NIH) macro and subsequently calculated using an established relationship as previously described ^64^.

### Computational fluid transport model for morphological analysis and velocity, flow rate, and WSS calculations

The micro-Vascular Evaluation System (μVES) ^65^ was used as the CFD model. μVES was used to calculate the flow rate, velocity, and WSS across every individual MVN segment. Flow rate boundary conditions were determined based on the experimentally measured MVN hydraulic resistances and corresponding flow rates. Maximum projected, 2D thresholded images were used as input to μVES. A custom vessel perturbation algorithm was added to μVES and used to conduct a vessel radius perturbation analysis. For the vessel perturbation study, the diameter of one vessel segment of interest was decreased by 10%, the complete MVN flow simulation was re-run using the same boundary conditions, and the WSS, velocity, flow rate, and diameter of the perturbed vessel segment was noted. This same procedure was done for each vessel segment of interest, perturbing one vessel segment at a time while keeping the diameters of the remaining MVN unperturbed. The derivative of the flow property (ie. WSS, flow rate, or velocity) with respect to the derivate of the radius was calculated as

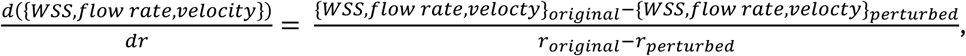

where *r* is the vessel radius.

### Inflammatory cytokine analysis

The full volume of conditioned media (200 µL) was collected from static devices on day 7 and from long-term flow devices on day 14 and 22 and used in a Human Focused 15-Plex Discovery Assay® (Eve Technologies). For all conditions, the medium was in contact with the MVNs for 24 hours before sampling. Samples of growth medium and maintenance medium were also analyzed for background signal, which was subtracted from all conditioned medium samples. The static devices were then connected to the pump for long-term flow, as previously described. Fresh media was added to the long-term flow devices and pump flow was resumed.

### Bulk RNA sequencing

For each timepoint analyzed, the fibrin hydrogel containing the MVNs was cut out of the devices and placed in 5 mL of 100 μg mL^−1^ Liberase (Millipore Sigma), 100 μg mL^−1^ DNase I (Sigma Aldrich), and 9.2 μg mL-1 Elastase (Thermo Fischer Scientific) in DMEM (Thermo Fischer Scientific) for 75 minutes at 37°C with intermittent agitation. Cells from 3-4 devices were pooled for each replicate. Dead cells were removed using the Dead Cell Removal Kit and an OctoMACS Separator (Miltenyi Biotec), cells were incubated with CD31 MicroBeads (CD31 MicroBead Kit, Miltenyi Biotec) and HUVECs and FBs were sorted using the OctoMACS Separator (Miltenyi Biotec). The sorted cells were lysed with TRIzol (Thermo Fischer) and stored at -80°C. Once the cell lysates from all timepoints were collected, the lysates were thawed, and 0.2 mL of chloroform (Millipore Sigma) per 1 mL of TRIzol reagent was added, the samples were centrifuged for 15 minutes at 12,000 x g at 4°C, and the aqueous phase was collected. The samples were then prepared using the “Preparation of Cell Pellets” protocol for RNA extraction using the chemagic 360 (Revvity). RNA-seq data analysis was conducted using a robust R-based pipeline. BAM files were quantified with featureCounts^66^ to generate a counts matrix, which was processed with DESeq2^67^ for differential gene expression analysis. Genes with low expression were filtered out, and more than four biological replicates per condition ensured statistical rigor. PCA confirmed sample clustering by condition. Pathway enrichment analysis was performed using clusterProfiler^68^, identifying significantly enriched pathways that contextualize the observed gene expression changes. The R code used for analyzing the counts files has been uploaded to the repository.

### Statistics

Average values are reported as mean ± standard deviation and sample sizes are reported in figure legends. Statistical analysis was performed using GraphPad Prism software, version 10. Parametric, two-tailed t tests were applied to analyses involving two groups and ordinary one-way ANOVA with Tukey’s multiple comparison tests were applied to analyses involving more than two groups. Experiments consisting of tracking the same MVNs over time used paired statistics. For the 2D analysis group separation analysis, the Henze-Zirkler test statistic was used to test for normality and a 2D permutation test with 10,000 permutations was used to as the test statistic. The random seed was set to 101 for all groups compared. A 2D group separation analysis using linear discriminant analysis was used to find a linear decision boundary between two groups in the 2D comparison. * indicates p<0.05, ** indicates p<0.01, *** indicates p<0.001, **** indicates p<0.0001, and n.s. indicates p>0.05.

## Author Contributions

MF performed all experiments and analyzed the data. MF, AB, GO designed and performed conceptual flow experiments. MF, EC, MFC, ZW designed the long-term flow experiments and contributed to the writing and editing of the manuscript. VV and AK performed the RNAseq analysis. All authors reviewed and/or edited the manuscript before submission.

## Acknowledgments

The authors are thankful to Charlie Demurjian for preparing the data for open access publishing, Stuart Levine for guidance in designing and executing experiments for bulk RNA sequencing, Luca Possenti and Alberto Rota for troubleshooting the μVES code. MF was supported by an MIT MathWorks Fellowship. EC was supported by an Early Postdoc.Mobility fellowship from the Swiss National Science Foundation (P2EZP2_199914), a postdoctoral fellowship from the Ludwig Center at MIT Koch Institute for Integrative Cancer Research, and a Postdoc.Mobility fellowship from the Swiss National Science Foundation (P500PB_222131). This work was supported by a grant from the National Cancer Institute (U54-CA261694). The funder played no role in study design, data collection, analysis and interpretation of data, or the writing of this manuscript.

