## Supplemental Information for "Remodeling of self-assembled microvascular networks under long term flow"

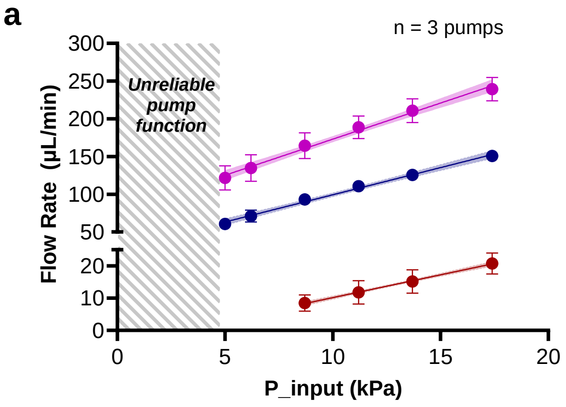


| | 1e11 Resistance | 1e12 Resistance | 1e13 Resistance |
|---|---|---|---|
| Equation | $y = 9.571x + 77.34$ | $y = 7.208x + 27.22$ | $y = 1.413x - 3.964$ |
| $R^2$ | 0.9928 | 0.9951 | 0.9988 |

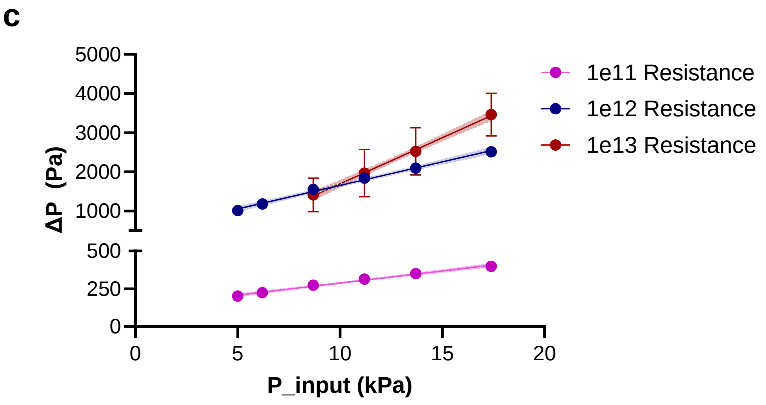


| | 1e11 Resistance | 1e12 Resistance | 1e13 Resistance |
|---|---|---|---|
| Equation | $y = 15.94x + 128.9$ | $y = 120.2x + 452.6$ | $y = 235.2x - 657.5$ |
| $R^2$ | 0.9927 | 0.9951 | 0.9988 |

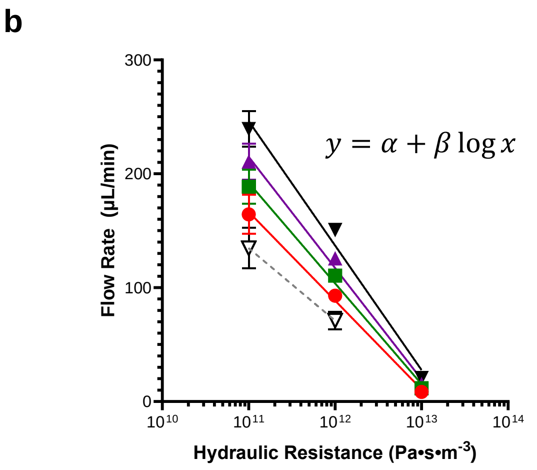


| P_input (kPa) | $\alpha$ | $\beta$ | $R^2$ |
|---|---|---|---|
| 8.7 | 1024 | -77.93 | 0.9801 |
| 11.2 | 1165 | -88.43 | 0.9840 |
| 13.7 | 1280 | -97.75 | 0.9847 |
| 17.4 | 1448 | -109.3 | 0.9808 |

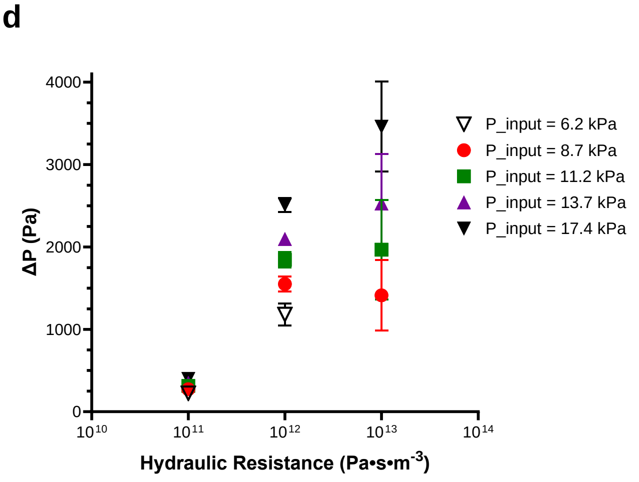


**Supp. Fig. 1. MicroHeart is characterized for a variety of hydraulic resistances.** (a-b) The pump flow rate was measured for varying input pressures and hydraulic resistances. The flow rate is plotted as a function of input pressure (a) and hydraulic resistance (b). The linear fits are indicated in the figures. The corresponding pressure difference was calculated and plotted as a function of input pressure (c) and hydraulic resistance (d). n = 3 pumps.

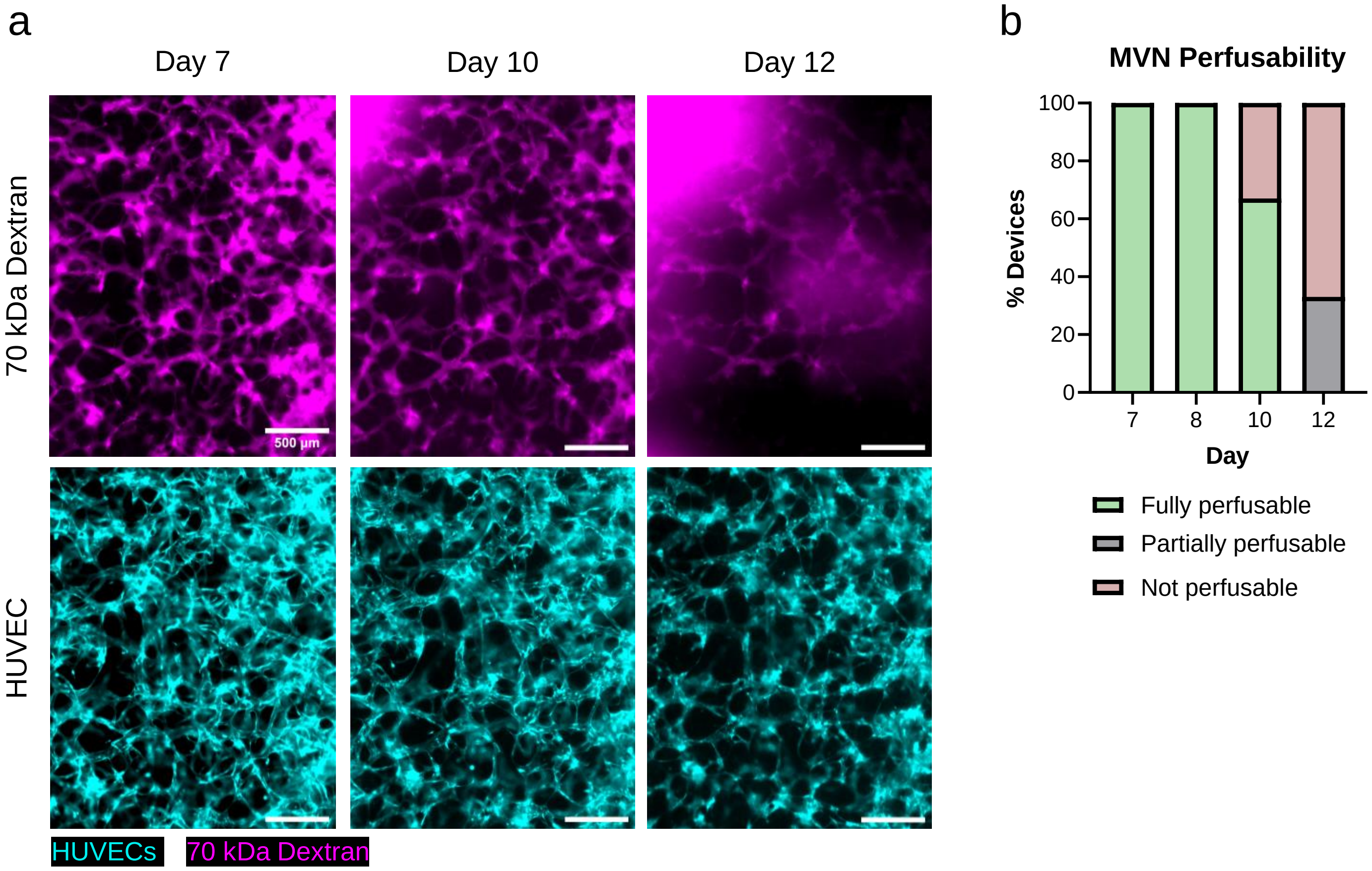


**Supp. Fig. 2. MVNs cultured with maintenance medium under static conditions regress and lose perfusion by day 12.** (a) Representative images of the same ROI over time. Scale bar is 500 μm. (b) The perfusability of MVNs is characterized over time. n = 3 MVNs.

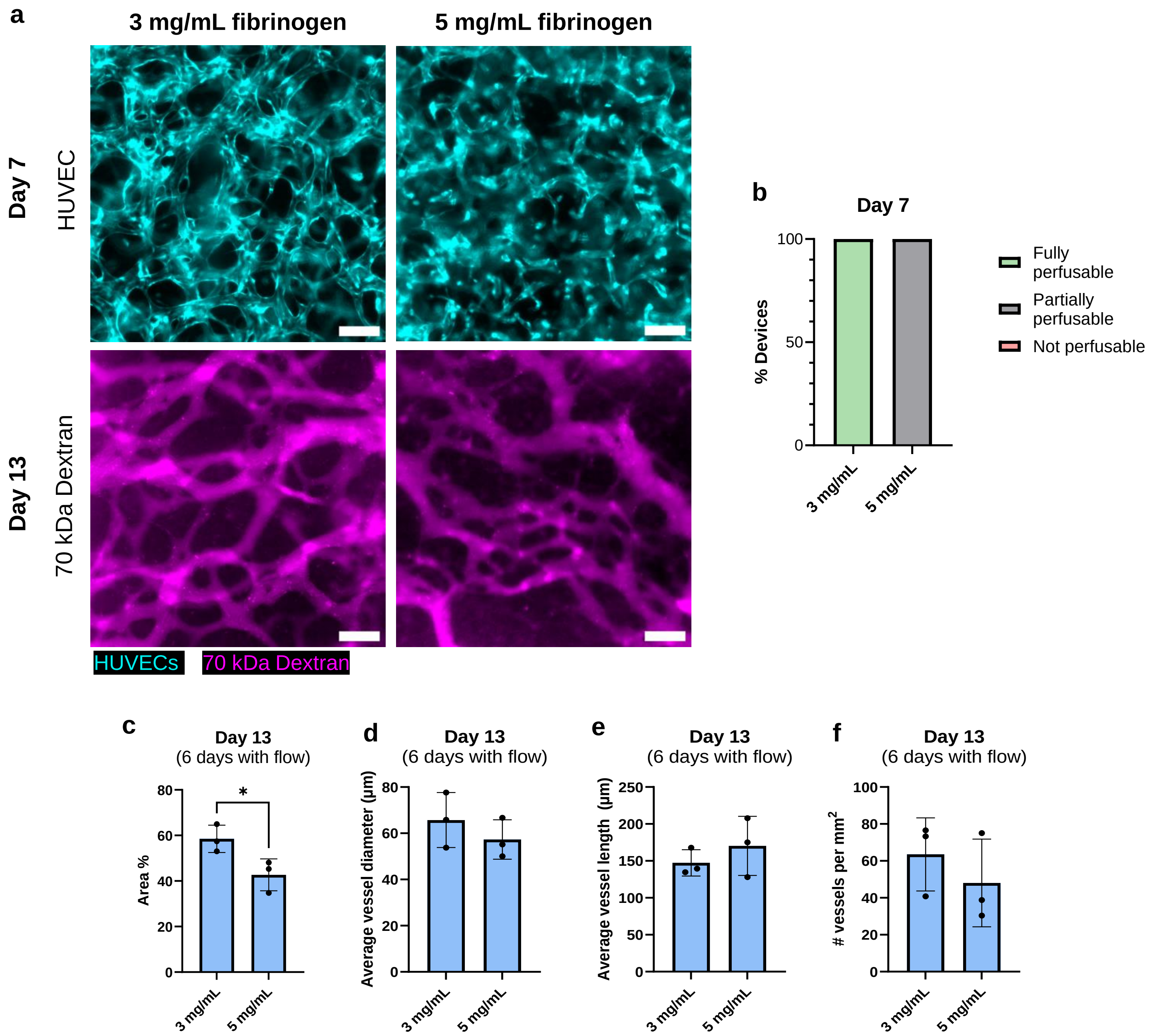


**Supp. Fig. 3. Pump flow yields similar vascular morphology after 6 days of flow despite different initial seeding conditions.** (a) Representative images of MVNs on day 7 and on day 13, after 6 days of pump flow. Scale bar is 200 μm. (b) Perfusability of the MVNs on day 7. (c) Vessel area %, (d) average vessel diameter, (e) average vessel length, and (f) vessel density are reported. n = 3 MVNs.

**Assumptions**

- Fully developed viscous flow
- Incompressible, Newtonian fluid

**Governing relationships**

1) Flow rate: $\frac{\rho g(z_1 - z_2)}{R} = Q$, where $z_1$ and $z_2$ are a function of time, $t$.
2) Mass conservation: $A_1\dot{z}_1 = -A_2\dot{z}_2$
3) Define Q: $Q = A_2\dot{z}_2$
4) Define $V_t$: $V_t = A_1z_1 + A_2z_2$

**Finding an expression for hydraulic resistance, *R***

We first begin with $$A_1\dot{z}_1 = -Q = \frac{-\rho g(z_1 - z_2)}{R}$$

Rearranging and substituting $V_t$ we get $$\dot{z}_1 = \frac{\rho g V_t}{RA_1A_2} - z_1\left(1 + \frac{A_1}{A_2}\right)\left(\frac{\rho g}{RA_1}\right)$$

Using the initial condition, $z_1(t = 0) = z_{1,0}$, where $z_{1,0}$ is a known and measured quantity, we can solve to get the following expression:

$$z_1(t) = \frac{V_t}{A_1 + A_2} + \left(z_{1,0} - \frac{V_t}{A_1 + A_2}\right)e^{\frac{-\rho g}{A_1R}\left(1+\frac{A_1}{A_2}\right)t}$$

Rearrange and take the natural logarithm to get

$$\ln\left(\frac{z_1(t) - \frac{V_t}{A_1 + A_2}}{z_{1,0} - \frac{V_t}{A_1 + A_2}}\right) = \frac{-\rho g}{A_1R}\left(1 + \frac{A_1}{A_2}\right)t$$

Using experimentally measured values for $z_1$ and $t$, find the slope, $m$, of the log term with respect to time. Use the following expression to plug in the slope and solve for the hydraulic resistance, $R$:

$$R = \left(1 + \frac{A_1}{A_2}\right)\left(\frac{\rho g}{A_1}\right)\frac{1}{m}$$

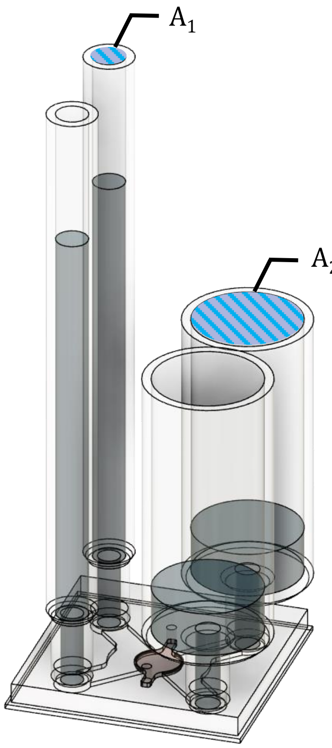


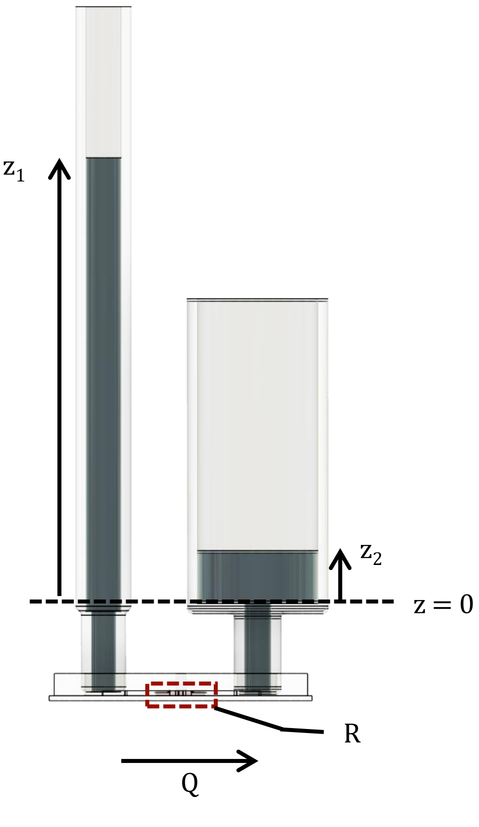

**Supp. Fig. 4. Analytical derivation of the relationship of the height of a hydrostatic pressure head with time for the case when the syringe diameter on the high-pressure side of the gel channel is not equal to the syringe diameter on the low-pressure side of the gel channel.**

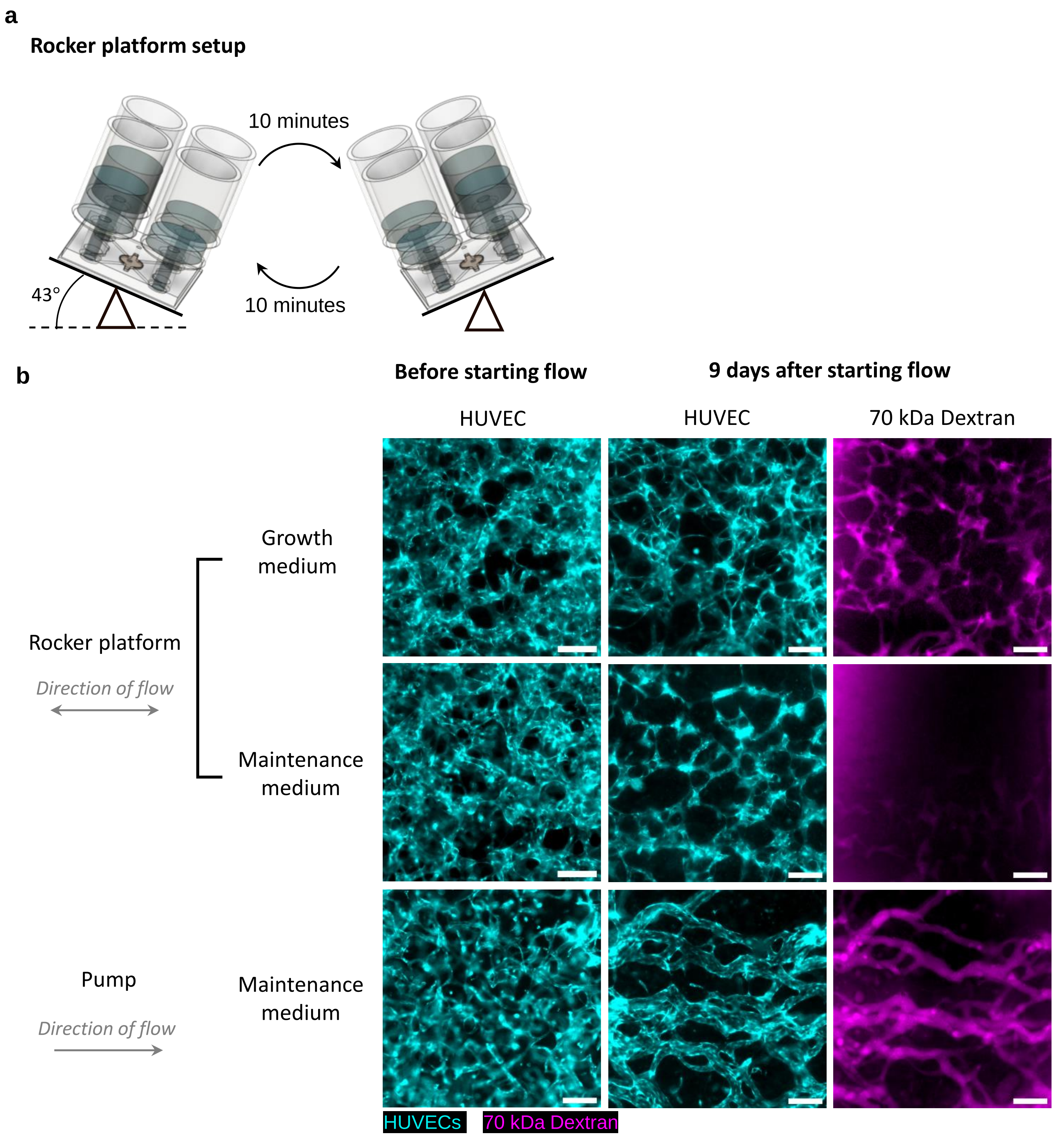


**Supp. Fig. 5. MVNs remodel differently depending on the medium and the method of perfusion used.** (a) Experimental setup for the rocker platform. (b) Representative images of MVNs at day 7 and at day 16 (9 days of flow). After images were taken at day 7, the MVNs were perfused using one of three methods: maintenance medium and rocker, growth medium and rocker, maintenance medium and pump. The direction of flow is indicated in the figure. Scale bar is 300 μm.

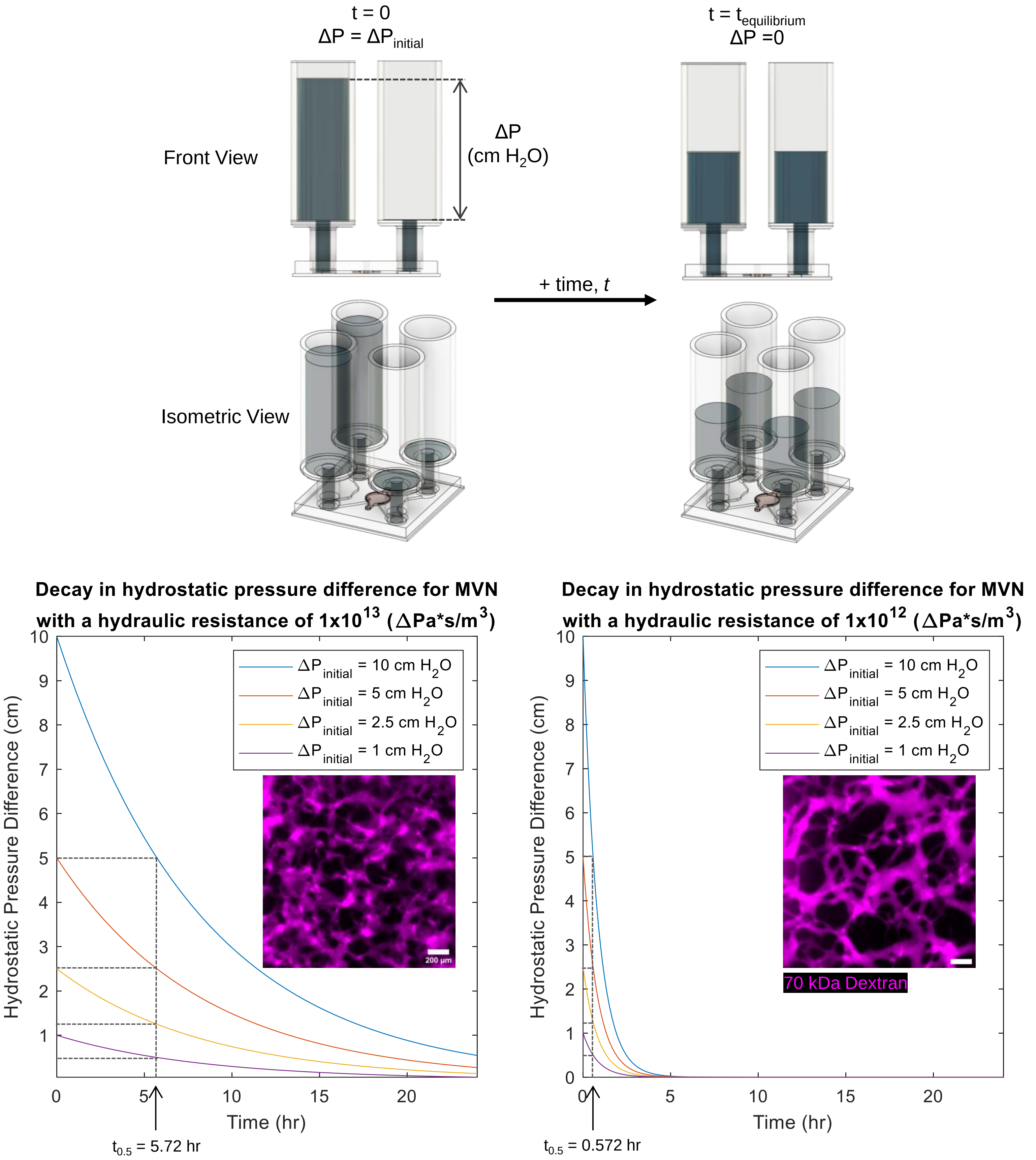


**Supp Fig. 6. Simulating the decay in hydrostatic pressure difference across a vascular bed based on an analytical relationship.** (a) The experimental setup used for the simulations consists of an initial hydrostatic pressure difference across a vascular bed at time, $t$ = 0. At a later time, $t_{equilibrium}$, the hydrostatic pressure difference equilibrates to zero. (b-c)The exponential hydrostatic pressure difference decay depends on the hydraulic resistance of the vascular bed. The half-lives of the hydrostatic pressure differences are labeled in the figures. Inset images show vascular beds characteristic of the hydrostatic resistance used for the simulation. Scale bar is 200 $\mu$m.

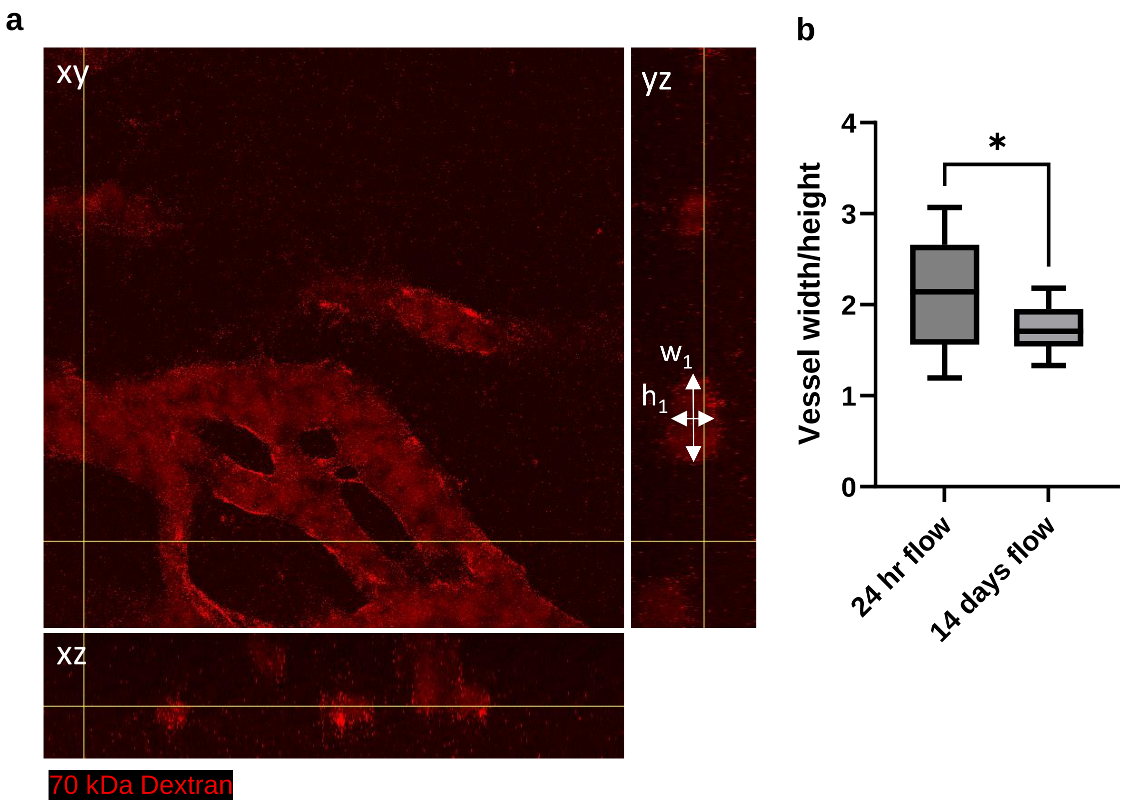


**Supp Fig. 7. The cross section of the MVNs is elliptical.** (a) Representative orthogonal image of a vessel cultured with 14 days of flow. (b) The aspect ratio of the major to minor axis of the cross section of a vessel decreases with 14 days of flow but remains elliptical. n = 19 across 4-5 devices.
